# Citrate cross-feeding between *Pseudomonas aerguinosa* genotypes supports *lasR* mutant fitness

**DOI:** 10.1101/2023.05.30.542962

**Authors:** Dallas L. Mould, Carson E. Finger, Nico Botelho, Stacie E. Stuut, Deborah A. Hogan

**Affiliations:** Geisel School of Medicine at Dartmouth, Department of Microbiology and Immunology, Hanover, NH USA

**Keywords:** quorum sensing, citrate, catabolite repression, cross feeding, LasR, CbrA, CbrB, RhlR, Crc

## Abstract

Across the tree of life, clonal populations—from cancer to chronic bacterial infections — frequently give rise to subpopulations with different metabolic phenotypes. Metabolic exchange or cross-feeding between subpopulations can have profound effects on both cell phenotypes and population-level behavior. In *Pseudomonas aeruginosa*, subpopulations with loss-of-function mutations in the *lasR* gene are common. Though LasR is often described for its role in density-dependent virulence factor expression, interactions between genotypes suggest potential metabolic differences. The specific metabolic pathways and regulatory genetics enabling such interactions were previously undescribed. Here, we performed an unbiased metabolomics analysis that revealed broad differences in intracellular metabolomes, including higher levels of intracellular citrate in LasR- strains. We found that while both strains secreted citrate, only LasR- strains, consumed citrate in rich media. Elevated activity of the CbrAB two component system which relieves carbon catabolite repression enabled citrate uptake. Within mixed genotype communities, we found that the citrate responsive two component system TctED and its gene targets OpdH (porin) and TctABC (transporter) required for citrate uptake were induced and required for enhanced RhlR signalling and virulence factor expression in LasR- strains. Enhanced citrate uptake by LasR- strains eliminates differences in RhlR activity between LasR+ and LasR- strains thereby circumventing the sensitivity of LasR- strains to quorum sensing controlled exoproducts. Citrate cross feeding also induces pyocyanin production in LasR- strains co-cultured with *Staphylococcus aureus*, another species known to secrete biologically-active concentrations of citrate. Metabolite cross feeding may play unrecognized roles in competitive fitness and virulence outcomes when different cell types are together.

**IMPORTANCE:** Cross-feeding can change community composition, structure and function. Though cross-feeding has predominantly focused on interactions between species, here we unravel a cross-feeding mechanism between frequently co-observed isolate genotypes of *Pseudomonas aeruginosa*. Here we illustrate an example of how such clonally-derived metabolic diversity enables intraspecies cross-feeding. Citrate, a metabolite released by many cells including *P. aeruginosa*, was differentially consumed between genotypes, and this cross-feeding induced virulence factor expression and fitness in genotypes associated with worse disease.

## INTRODUCTION

In populations formed by diverse cell types, from microbes to human cells, *de novo* mutation and selection can lead to subpopulations with different metabolic phenotypes (1, 2, 3). Genotypic and metabolic heterogeneity can make populations more resilient but can also hinder successful eradication of populations in the context of disease (4, 5, 6, 7).

Studies on the diversity of *P. aeruginosa* isolates from chronic infections find mutations in the same genes or pathways, and often single infections contain genetically diverse populations. *P. aeruginosa* isolates from later stage disease often exhibit amino acid auxotrophies, altered iron acquisition strategies, altered exopolysaccharide production, and reduced activity of a major microbial communication system referred to as quorum sensing (QS) (8). A reduction in QS activity in *P. aeruginosa* isolates most often results from loss-of-function mutations in the gene encoding the central regulator LasR, while the RhlR- and PqsR-dependent transcriptional networks of QS in *P. aeruginosa* isolates remain comparatively conserved (9). Interestingly, *lasR* mutants also readily arise in cultures *in vitro* under different experimental evolution regimes suggesting that loss of LasR function may provide a general fitness advantage even outside the contexts of disease (10, 11, 12).

In addition to its role in density-dependent activation of secreted virulence factors, QS control of metabolism has also been broadly observed (13, 14, 15). In *P. aeruginosa*, differences in metabolism explain in part the rise of the commonly observed *lasR* mutants (16, 17, 18). Experimental evolution studies show that the rise of LasR- strains is affected by the nutritional environment (11) and a metabolic regulatory process known as carbon catabolite repression (18). Carbon catabolite repression (or catabolite repression) controls carbon utilization through a post-transcriptional process wherein the co-repressor Crc blocks translation of transcripts for alternative carbon utilization when a preferred substrate is present (19, 20). Activity of the CbrAB two component system relieves Crc-dependent translational repression (21) and is required for the reproducible rise of LasR- strains *in vitro* (18). The increased CbrAB activity in LasR- strains enables enhanced consumption of less preferred carbon sources, including the metabolites enriched in progressive cystic fibrosis (CF) airways from where LasR- strains are frequently isolated (18, 16). The commonly observed genetic changes that influence metabolism and the production of QS-controlled goods can alter the host environment as well as the interactions among neighboring host and microbial cells, including kin.

LasR- strains interact differently with host cells and other microbes than their LasR+ counterparts (22, 23, 24, 25), and these differences may help explain why LasR- strains are associated with worse health outcomes (26) yet express fewer virulence factors in monoculture and show low virulence in model systems of infection (27, 28, 29, 30). This apparent contradiction may extend from their altered metabolism (16, 18), differential immune responses (23, 24), or the diversification noted *in vivo* where different genotypes often coincide (31, 26). Though *lasR* mutants are common, they are frequently co-isolated with LasR+ strains (8). For example, analysis by Hoffman et al. of multiple isolates from 58 different people with CF revealed twenty individuals had infections that contained both wild-type and *lasR* mutant cells (26). Mixed communities containing both LasR+ and LasR- strains produce more secreted virulence factors than either strain grown in isolation (32). Surprisingly, given the positive role LasR plays in the regulation of many secreted factors, LasR- strains are responsible for the hyper-production of the antagonistic factors in co-cultures with LasR+ cells (32). Because *lasR* mutants are associated with worse disease outcomes (25, 26), it is important to under-stand both the factors contributing to their rise as well as the consequences once they emerge. Here we characterize the metabolic underpinnings that enable cross-feeding between common *P. aeruginosa* variants and illustrate its protective role in microbial fitness.

## RESULTS

### LasR has broad impacts on the metabolome including effects on intracellular and extracellular citrate concentrations

To investigate the impact of LasR on *P. aeruginosa* metabolism, we performed metabolomics analysis on colony biofilms of *P. aeruginosa* strain PA14 wild type and its Δ*lasR* derivative and a pair of clinical isolates that differed by LasR function (DH2417 LasR+ and DH2415 LasR-). Each strain was grown in quintuplicate on both Lysogeny Broth (LB) and Artificial Sputum Medium (ASM) (33, 34), which mimics the CF lung environment. Analysis of the intracellular metabolomes, performed by ultrahigh performance liquid chromatography-tandem mass spectroscopy (UPLC-MS/MS), revealed broad differences between LasR+ and LasR- cells. Of the 562 detected metabolites with known identities, over 100 were significantly differentially abundant in each comparison: Δ*lasR* / WT on LB, DH2415 (LasR-) / DH2417 (LasR+) on LB and Δ*lasR* / WT on ASM (Table S1). In a principal component analysis (PCA) of the normalized metabolite counts, the first component, which explained 47.38% of the variation in the data, separated samples by medium type while the second component, explaining 21.02% of the variation in the data, appeared to separate samples by LasR functional status (Fig. 1A). The clinical isolates clustered more closely to the Δ*lasR* strain than the wild type on LB, but still showed separation by LasR type along the second principal component axis (Fig. 1A).

**FIG 1.**
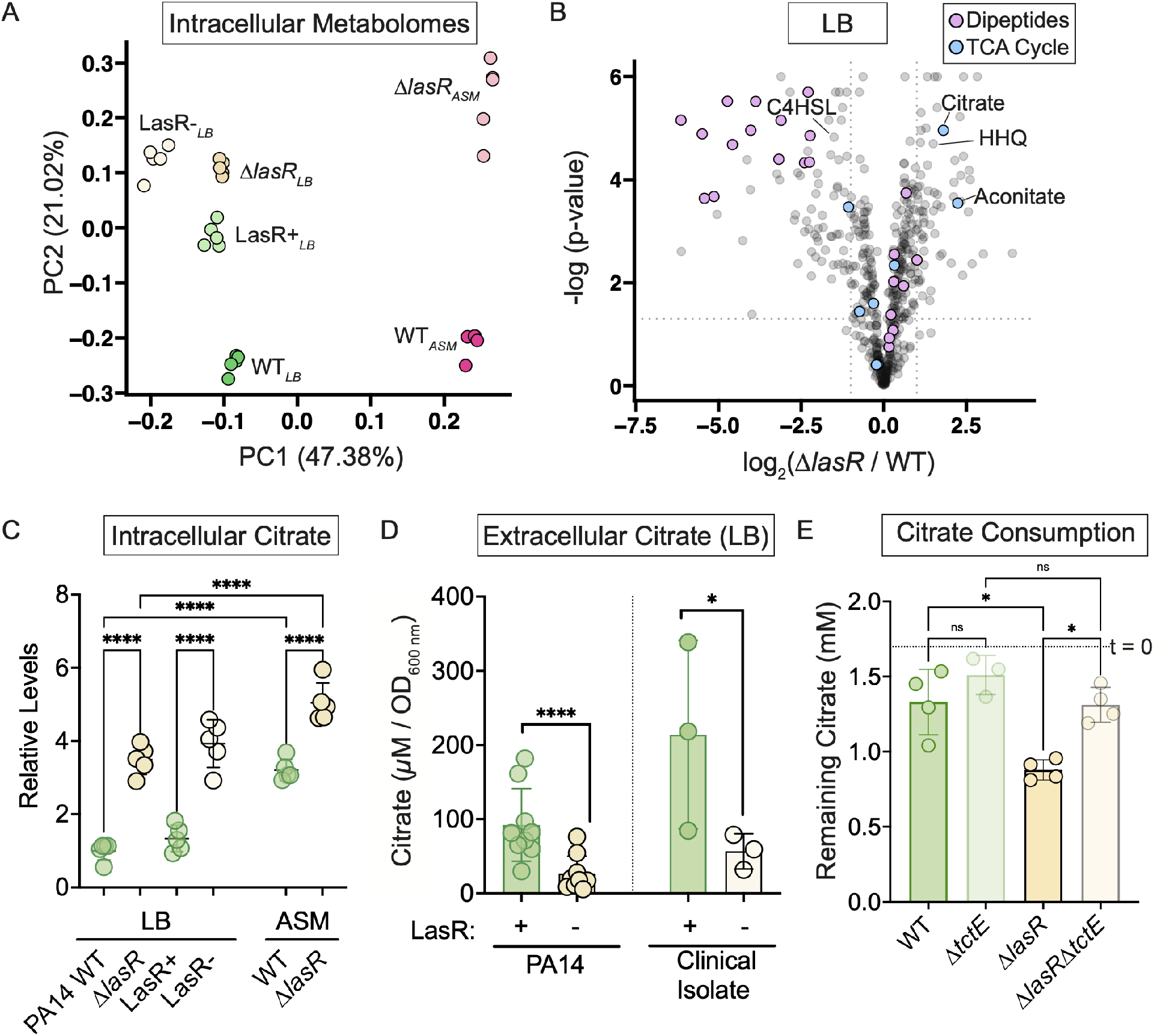
LasR function impacts the metabolic state of the cell promoting differences in citrate homeostasis. A. Principal component analysis of pareto normalized metabolite counts as determined by ultrahigh performance liquid chromatography-tandem mass spectroscopy (UPLC-MS/MS) of laboratory strains PA14 and Δ*lasR*, and closely-related CF clinical isolates with functional LasR (LasR+, DH2417) and dysfunctional LasR (LasR-, DH2415) grown as colony biofilms for 16 hours on LB and/or ASM with medium indicated as subscript. (n = 5) B. Volcano plot showing differential metabolite counts (log_2_ Fold Change) for Δ*lasR* cells relative to wild-type cells in LB on the x-axis; the y axis shows the −log_10_ (P value) for the difference between sample types. Metabolites categorized as dipeptides (lavender) or as part of the tricarboxylic acid (TCA) cycle (light blue) are indicated alongside 4-hydroxy-2-heptylquinoline (HHQ) and N-butanoyl-L-homoserine lactone (C4HSL) which are known to be enriched and depleted in *lasR* mutants, respectively. C. Relative metabolite counts of citrate as determined from experiment described in A for the laboratory strain on LB and ASM and the clinical isolate pair on LB. Statistical significance determined by Ordinary One-way ANOVA with Šidák’s multiple comparison’s test: ****, P value < 0.001. D. Extracellular citrate measured in supernatant of PA14 (n = 9), Δ*lasR* (n = 9), DH2417 (LasR+) (n = 3), and DH2415 (LasR-) (n = 3) of 24 h grown cultures in 5 mL of LB using an enzymatic commercial assay. Statistical significance determined by Student’s t-test, ****, P value < and *, P value < 0.05. E. Citrate remaining in supernatants taken from densely inoculated cultures of wild type, the Δ*lasR* strain, and derivatives lacking the citrate-responsive sensor kinase TctE incubated for 24 h in LB supplemented with 2 mM citrate as measured via enzymatic assay. (n = 4) Dotted line indicates citrate concentration recovered at the start of the experiment (t = 0). Data included in this graph are a subset of the experiment including data in Fig. 2C. Statistical significance determined via Ordinary One-Way ANOVA with Sidaks multiple comparison’s test for entire experimental datasets: **, P value < 0.005; ns, not significant (P value = 0.75 and 0.66 for WT vs. Δ*tctE* and Δ*tctE* vs. Δ*lasRΔtctE* comparisons, respectively.)

The metabolomics data were consistent with well-described roles for LasR in the control over the RhlR- and PqsR (MvfR)-dependent regulons of the inter-connected quorum sensing system and the positive regulation of protease production. For example, LasR regulates the enzyme PqsH that converts 4-hydroxy-2-heptylquinoline (HHQ) into 2-heptyl-3-hydroxy-4(1H)-quinolone (PQS), and loss of Δ*lasR* leads to the accumulation of HHQ which gives Δ*lasR* mutant colonies their characteristic sheen or metallic appearance (16). We found that HHQ was significantly more abundant in LasR-cells when compared to the wild type on LB (Fig. 1B) and ASM in both the PA14 strain and clinical isolate backgrounds to a lesser extent (Fig. S1A & B). Also, consistent with LasR-dependent production of the autoinducer N-butanoyl-L-homoserine lactone (C4HSL) synthase RhlI (regulated by RhlR), we found significantly lower levels of C4HSL in LasR-cells relative to wild type on LB (Fig. 1B). Of note, unlike the LB samples (Fig. 1B & Fig. S1B), C4HSL was not significantly differentially abundant between strains on ASM (Fig. S1A). Lastly, over 30% of all dipeptides detected were significantly less abundant in Δ*lasR* cells consistent with the lower proteolytic activity present in LasR- strains (35, 36) (Fig. 1B). Similar observations were made when comparing the wild type and Δ*lasR* strain on ASM and the LasR+ and LasR-clinical isolates on LB (Fig. S1A & B).

The broad differences in the metabolite profiles between LasR+ and LasR-cells suggest different anabolic and catabolic states when grown on the same medium type which creates the potential for cross-feeding or differential substrate utilization between strains. Tricarboxylic acid (TCA) cycle intermediates are critical for both energy generation and biosynthesis, and are frequently secreted by microbes and host cells under particular conditions like iron or oxygen limitation (37). We found that in both strain backgrounds on LB and in strain PA14 on ASM, one intermediate in the TCA cycle, citrate, stood out with a fold change > 1.6 and P value < 0.05 in all conditions (Fig. S1C). Succinate was also significantly different in all conditions, but the fold differences were much smaller. Intracellular citrate was significantly more abundant in LasR-cells than those with intact LasR on LB (Fig. 1C). Given our prior work that found that citrate supplementation induces RhlR-dependent quorum sensing activity in LasR- strains but not LasR+ strains (32), we focused our efforts on citrate with the goal of understanding if it is managed differently in LasR+ and LasR-cells.

To complement the analysis of intracellular citrate, we examined extracellular citrate levels in cell-free supernatants from the same sets of LasR+ and LasR-isolates as above after 16 h of growth in LB medium. In fresh, uninoculated LB medium we detected 100 ± 103 *μ*M (average± s.d.) citrate (Fig. S2). After growth, PA14 wild-type culture supernatants contained 327± 89 *μ*M citrate while supernatants of Δ*lasR* cultures contained only 87 ± 58 *μ*M citrate (Fig. S2). The significantly higher levels of citrate in supernatants of wild-type cultures than in the LB medium blank suggested that wild-type cells secreted citrate. In contrast, there was no significant difference in citrate levels between the LB medium and supernatant from Δ*lasR* cultures (Fig. S2). The differences in supernatant citrate were not due to differences in growth as OD-normalized citrate levels were significantly lower (by 3- to 4-fold) in the supernatants of LasR- strains when compared to those for their respective LasR+ strains (Fig. 1D). Overall, relative to LasR+ cells, LasR-cells had higher intracellular citrate levels and less extracellular citrate to suggest LasR function may impact citrate consumption and/or secretion.

To monitor citrate consumption, we added 2 mM citrate to high density cultures in LB medium and monitored the amount of citrate remaining after 24 hours. As a control, we included strains lacking the gene encoding the histidine kinase TctE, which regulates genes involved in citrate uptake (Fig. S3A) (38, 39, 40). In the supernatant of the Δ*lasR* strain, there was significantly less citrate recovered than for the wild type (Fig. 1E). The depletion of citrate by LasR- strains required *tctE*, and there was no significant difference between the Δ*tctE* and Δ*lasR*Δ*tctE* strains (Fig. 1E). There was no significant difference in the amount of citrate remaining in the supernatant of PA14 wild type compared to supernatant from the Δ*tctE* strain to suggest minimal net citrate consumption by the wild type (Fig. 1E). Collectively, the data suggest differential consumption by LasR+ and LasR- strains of at least one metabolite: citrate.

### Reduced catabolite repression via CbrB-Crc control promotes TctED-dependent citrate consumption by LasR- cells

LasR- strains have increased activity of the CbrAB two component system that enables de-repression of the catabolism of diverse substrates including citrate by modulating translational inhibition by Crc (16, 18) (Fig. 2A). Genetic analyses demonstrate the control of citrate catabolism by CbrAB and Crc in the Δ*lasR* mutant (Fig. 2B). Growth of the Δ*lasR* strain on citrate was significantly reduced in the absence of *cbrB*, and this could be complemented by restoration of *cbrB* at the native locus. Subsequent deletion of *crc* in the Δ*lasR*Δ*cbrB* strain also rescued growth on citrate and this phenotype was reversed by complementation of *crc* (Fig. 2B). The CbrB-Crc control of growth on citrate mirrored the effects by CbrB and Crc on citrate uptake by Δ*lasR* cells in LB (Fig. 2C). The Δ*lasR* mutant depleted more citrate than did the Δ*lasR*Δ*cbrB* strain, and again the Δ*lasR*Δ*cbrB* phenotype was relieved by deletion of *crc* (Fig. 2C). The Δ*lasRΔcbrB* defect in citrate depletion was similar to that observed for the Δ*lasRΔtctE* strain (Fig. 2C). Multiple lines of evidence indicate that the TctE-regulated genes *opdH-tctA-tctB-tctC* important for citrate utilization are under Crc control: (i) there is a putative *crc* binding site motif upstream of the translational start site (41); (ii) transcript levels of genes in this operon are elevated in *crc* mutants (41, 38), (iii) OpdH levels are elevated in strains lacking *crc* (42), and (iv) paired transcriptome and proteome analyses predict *tctC* and *opdH* as Crc targets (43).

**FIG 2.**
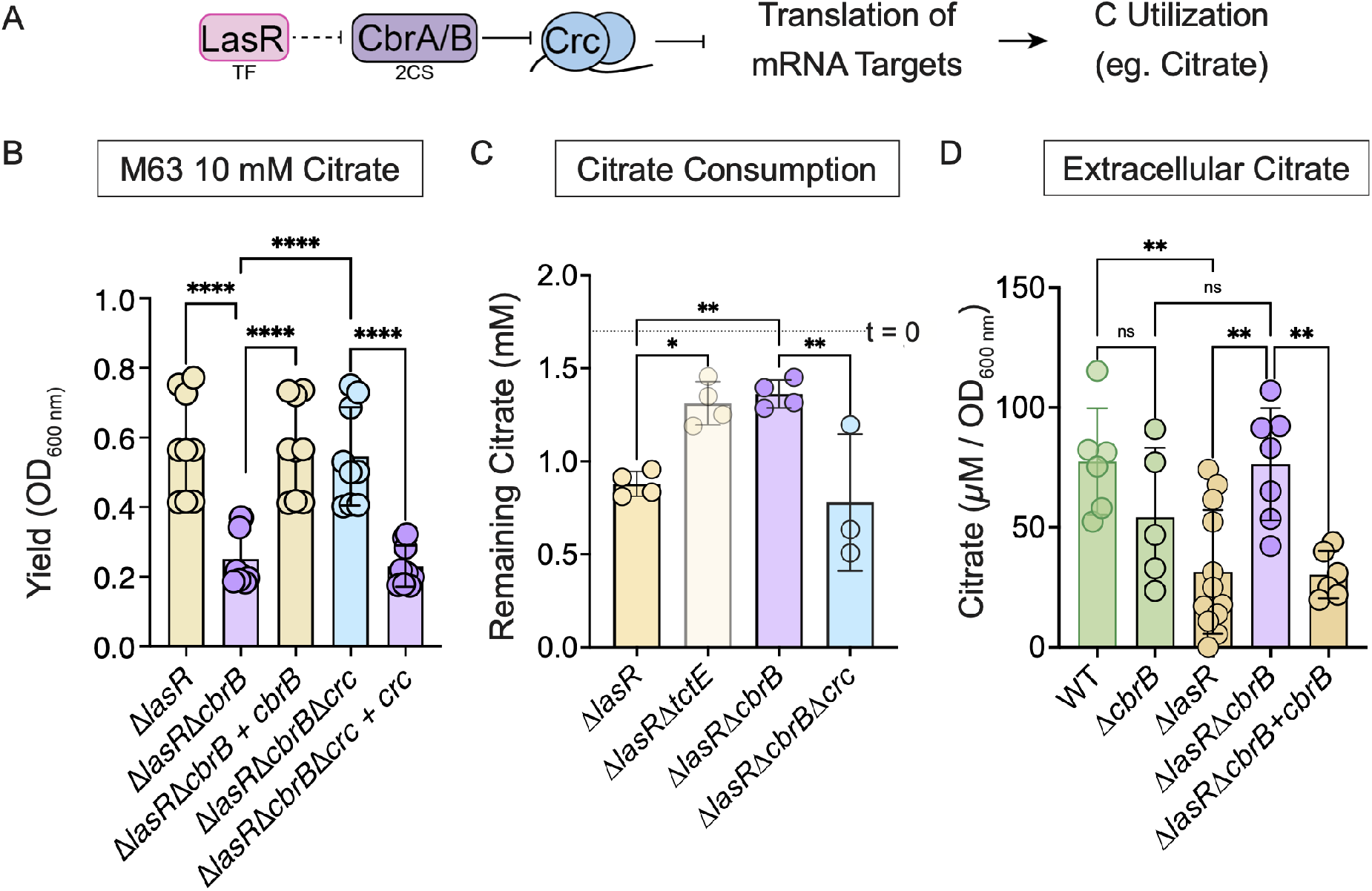
Catabolite repression controls citrate consumption but not citrate secretion enabling preferential uptake by LasR-cells. A. Simplified schematic illustrating the connection between the transcription factor (TF) LasR and the carbon catabolite repression pathway, including the two-component system (2CS) CbrAB which promotes de-repression of mRNA targets important for alternative carbon utilization such as citrate for which genes involved in its consumption are predicted to be under Crc translational inhibition. B. Growth of indicated strains on 10 mM citrate in M63 base after 16 hours following subculture from LB overnight. Data points indicate three biological replicates of three independent experiments. Statistical significance as determined by Ordinary One-Way ANOVA with Šidák’s multiple comparison’s test: ****, P value < 0.0001. C. Citrate consumption after 24 h incubation in LB supplemented with 2 mM citrate for the Δ*lasR* strain and derivatives lacking the sensor kinase TctE, the response regulator CbrB and/or the translational co-repressor Crc of the catabolite repression system. Each data point is from an independent experiment (n > 4). Dotted line indicates citrate concentration recovered at the start of the experiment (t = 0). Data included in this graph are a subset of the experiment including data in Figure 1E. Statistical significance determined via Ordinary One-Way ANOVA with Šidák’s multiple comparison’s test including all data from the experiment: *, P value = 0.02; **; P value < 0.01. D. Extracellular citrate levels (*μ*M / OD600 nm) in supernatants of the indicated strains grown in LB as measured via an enzymatic assay. Statistical significance as determined by Ordinary One-Way ANOVA with Sidak’s multiple comparison’s test: **, P value < 0.005; ns, not significant with P value > 0.44. (n > 5).

In order to determine if the differences in extracellular citrate were solely due to differences in citrate uptake or if the wild type and the Δ*lasR* mutant also differed in citrate production, we measured extracellular citrate in strains lacking citrate uptake due to an absence of *cbrB*. While the wild type had significantly more citrate than the Δ*lasR* strain, the levels of citrate were relatively high in both backgrounds when *cbrB* had been deleted (Fig. 2D). The loss of *cbrB* in the wild type did not significantly impact extracellular citrate levels, though the data may suggest that CbrB does play some condition-specific role in citrate secretion (Fig. 2D).

### In co-culture, tricarboxylates are detected by the TctED two component system of LasR- strains

In light of our data that suggested that *lasR* mutants are both producing and consuming citrate in LB medium, while the wild type cells only showed evidence for citrate release, we sought to determine if wild-type and Δ*lasR* strains exhibited similar transcriptional responses to citrate. To do so, we used a transcriptional reporter for the promoter upstream of the *opdH-tctA-tctB-tctC* operon which is induced in wild type (LasR+ backgrounds) by TctED in response to citrate (38, 40, 39). Consistent with the TctE-dependent growth (Fig. S3B) and consumption of citrate by *lasR* mutants (Fig. 1E), *tctE* and *tctD* both induced the *opdH* promoter in response to citrate in a LasR- strain background (Fig. 3A), much like reports in LasR+ strains (39). Complete induction of *opdH* promoter activity by citrate in *lasR* mutants required citrate entry into the cell as absence of the porin OpdH reduced induction by citrate, and absence of the inner membrane transporter TctABC rendered the *opdH* promoter completely non-responsive to citrate (Fig. S3C).

**FIG 3.**
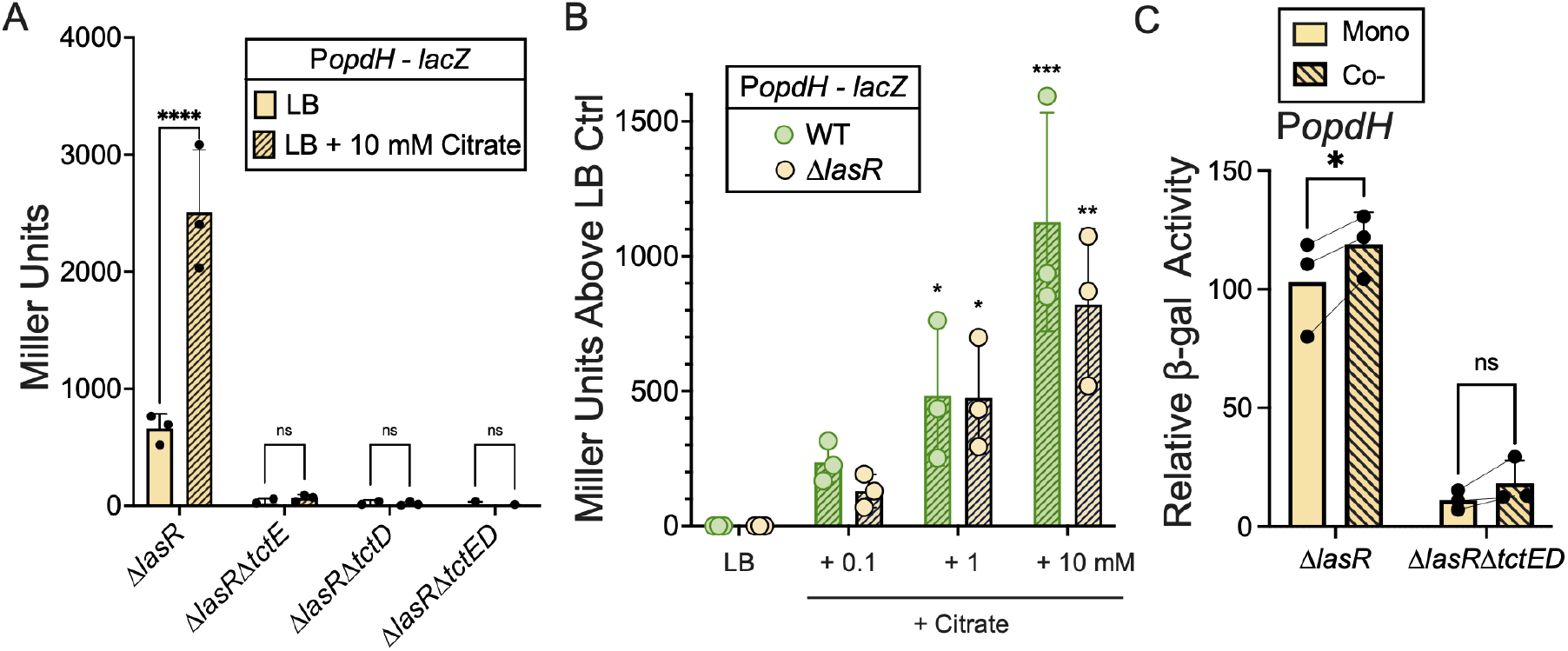
Citrate is detected in co-cultures of LasR+ and LasR- strains. A. Citrate responsive P*opdH*– *lacZ* activity quantified by Beta-galactosidase (*β*-gal) assay of Δ*lasR*, Δ*lasRΔtctE*, Δ*lasRΔtctD*, Δ*lasRΔtctED* colony biofilms on LB or LB supplemented with 10 mM citrate. B Promoter activity quantified for P*opdH* – GFP – *lacZ* of wild type and Δ*lasR* colony biofilms grown on LB supplemented with 100 *μ*M, 1 mM or 10 mM of citrate after 16 h of growth (n = 3) via beta-galactosidase (*β*-gal) assay. Statistical significance determined by Two-Way ANOVA (matching for citrate concentration and replicate) with Šidák’s multiple comparison’s test. There was no significant source of variation detected between strains (P value = 0.53), but there were differences noted across citrate concentrations (P value = 0.0004). Asterisks denote statistical significance for comparison to LB control: *, P value < 0.04; **, P value < 0.003; ***, P value = 0.0005. C. *β*-gal quantification of P*opdH* – GFP – *lacZ* in Δ*lasR* and Δ*lasRΔtctED* strains in monoculture or co-culture with fluorescently (mKate) labelled wild type (WT).

To assess the sensitivity of *opdH* promoter induction between strains, we considered concentrations within range of that observed in supernatant from wild type (LasR+) monocultures (∼200-500 *μ*M). Citrate induced P*opdH* - *lacZ* activity above baseline in both wild-type and Δ*lasR* cells when added to LB at low concentrations (+100 *μ*M) with more significant differences as citrate concentrations increased (Fig. 3B). Furthermore, we found significantly greater TctED-dependent induction of the *opdH* promoter fusion in Δ*lasR* cells when grown in co-culture with the wild type than when grown alone, suggesting Δ*lasR* cells were responding to the higher concentrations of extracellular citrate released by the wild type (Fig. 3C).

### CbrAB-Crc regulation is required for induction of RhlR signalling and RhlR-regulated pyocyanin production in LasR- strains grown in co-culture with LasR+ strains

We have previously reported that in LasR+/LasR-co-cultures, the presence of LasR+ cells induces RhlR activity in LasR-cells due to a factor other than acylhomoserine lactone autoinducers (32). Here, we sought to determine if citrate uptake was required for the induction of RhlR-dependent signaling and phenazine production in LasR- strains when cultured with their LasR+ counterparts. We monitored *rhlI* promoter activity in both single strain and mixed strain cultures. In co-culture, RhlR signaling in the Δ*lasR* strain increased when assessed at the community level with the beta-galactosidase assay (Fig. 4A), and the RhlR-dependent induction was validated at the cellular level via flow cytometry (Fig. S4). TctED and CbrB, which are required for citrate consumption, were also required for co-culture induced RhlR transcriptional activity in LasR-cells (Fig. 4A). We demonstrated that *rhlI* promoter activity increased in Δ*lasR* cells as citrate concentrations increased (Fig. 4B), and as expected, *rhlI* promoter activity was not observed in either the absence or presence of citrate in a Δ*lasR*Δ*rhlR* strain (Fig. 4B). The induction of *rhlI* promoter activity by citrate depended on TctED and one of its gene targets *opdH* involved in citrate uptake (Fig. 4C).

**FIG 4.**
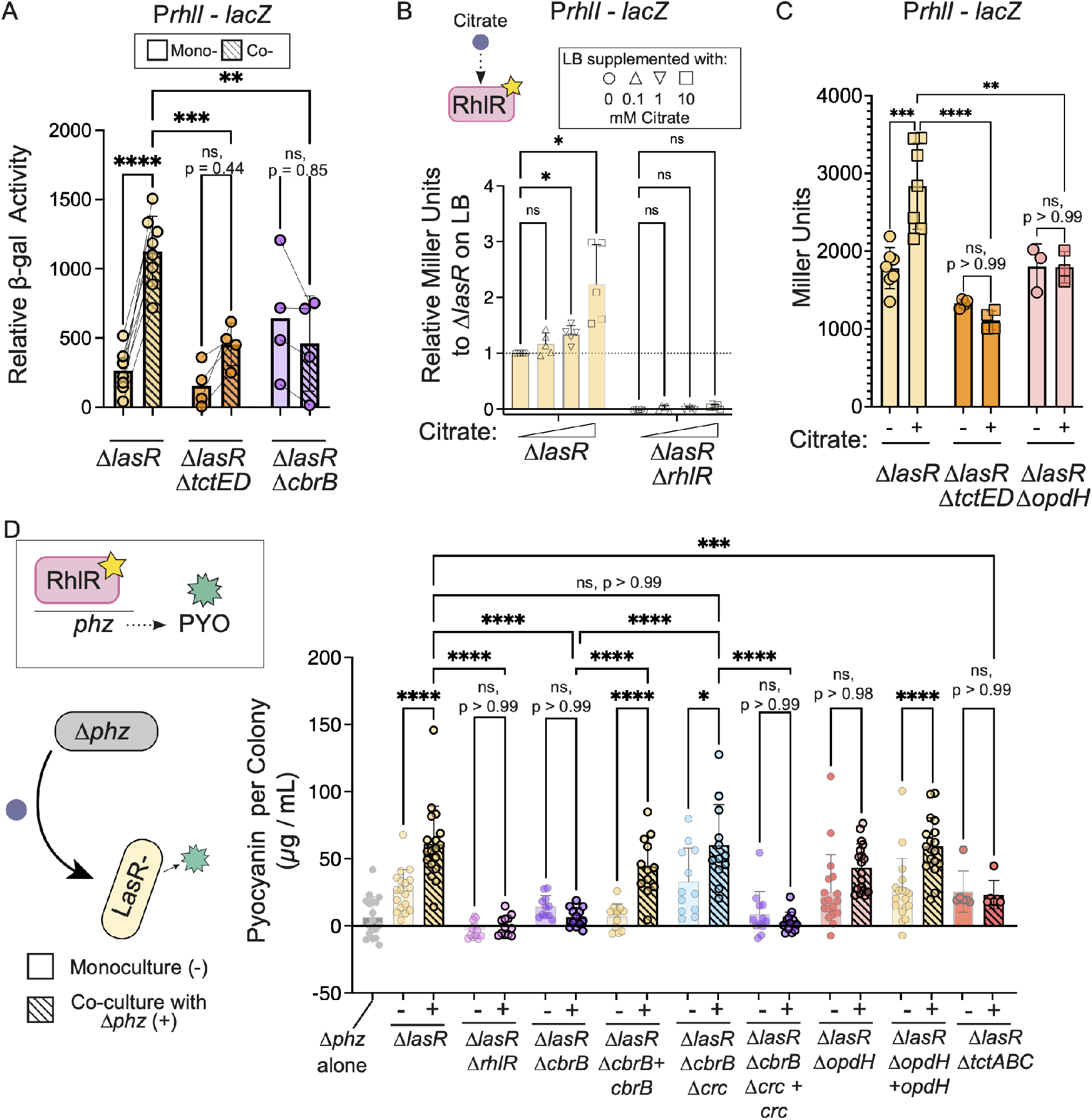
Redundant citrate uptake is required for activation of RhlR signalling and phenazine production by LasR- strains in co-culture. A. P*rhlI* – *lacZ* promoter activity quantified for Δ*lasR*, Δ*lasR*Δ*rhlR*, Δ*lasR*Δ*tctED*, and Δ*lasR*Δ*opdH* monocultures on LB agar supplemented with 0, 100 *μ*M, 1 mM, or 20 mM citrate. B. P*rhlI* – *lacZ* promoter activity for the indicated strains on LB agar (-) and LB agar supplemented with 20 mM citrate as quantified by Beta-galactosidase assay. C. Relative P*rhlI* – *lacZ* promoter activity quantified for the indicated *lasR* mutant derivatives in co-culture with fluorescently labelled WT (lacking a *lacZ* reporter). Miller units were adjusted for the proportion of the cellular density containing a strain with a *lacZ* reporter used as input in the *β*-galactosidase assay at the time of quantification. D. Pyocyanin is one product regulated by RhlR/I. Pyocyanin (PYO) levels (*μ*g / mL) per colony biofilm measured for monocultures (grey outline) of a strain lacking both phenazine biosynthesis operons (Δ*phz*,grey), the Δ*lasR* strain, and its mutant derivatives, as indicated, relative to co-cultures (black outline, hashed bars) of the Δ*lasR* mutants grown with the Δ*phz* strain. Data points are from at least three independent experiments with two biological replicates each. Statistical significance determined via Ordinary One-Way ANOVA with Šidák’s multiple comparison’s test. P values (p) as indicated.

RhlR regulates phenazine production, and thus we also monitored pyocyanin production by LasR- strains grown as colony biofilms in monoculture and co-culture with a LasR+ strain lacking both phenazine biosynthesis operons (Δ*phzA1*Δ*phzB1*Δ*phzC1*-*phzD1*Δ*phzE1*Δ*phzF1*Δ*phzG1*Δ*phzA2*Δ*phzB2*Δ*phzC2*Δ*phzD2*Δ*phzE2*Δ*phzF2*Δ*phzG2*) referred to as the “*phz*” strain. By co-culturing the *phz* strain with the Δ*lasR* strain and its derivatives, we are able to specifically measure LasR- strain phenazine production even in co-culture. Consistent with prior work (32), when the Δ*lasR* strain was co-cultured with the Δ*phz* strain (i.e. Δ*phz* / Δ*lasR*), there was a ~ 5-fold increase in pyocyanin compared to the Δ*lasR* monoculture (60 versus 14 *μ*g / mL of pyocyanin per colony biofilm) (Fig. 4D). Consistent with RhlR regulation of phenazine biosynthesis (32), no pyocyanin was recovered from the Δ*lasRΔrhlR* monoculture or co-culture with the Δ*phz* strain (Fig. 4D).

To determine if the increased activity of CbrB in LasR- strains was necessary for phenazine production, we grew the Δ*lasRΔcbrB* strain, which exhibits a more repressed metabolism, in monoculture and in co-culture with the Δ*phz* strain. Colony biofilms of the Δ*lasRΔcbrB* monoculture produced pyocyanin at concentrations nearly identical to that for Δ*lasR* monocultures (Fig. 4D). However, unlike the Δ*lasR* strain, pyocyanin production by the Δ*lasR*Δ*cbrB* mutant was not induced in the presence of the Δ*phz* strain and was significantly lower than when Δ*lasR* was co-cultured with the Δ*phz* strain (Fig. 4D). When the *cbrB* gene was complemented in the Δ*lasR*Δ*cbrB* strain (Δ*lasR*Δ*cbrB* + *cbrB*), co-culture pyocyanin was restored (Fig. 4D).

Furthermore, when the Δ*lasR*Δ*cbrB* strain was modified to delete *crc*, which encodes a co-repressor of the catabolite repression system that is downstream of CbrAB, co-culture pyocyanin production was also restored. While Δ*lasR*Δ*cbrB*Δ*crc* monocultures produced slightly higher pyocyanin levels than monocultures of Δ*lasR*, the levels increased from 29 *μ*g / mL in monoculture to 63 *μ*g /mL of pyocyanin in co-culture with the Δ*phz* strain (Fig. 4D). To ensure the restoration of the co-culture phenazine response was due to the loss of Crc-mediated repression, we complemented *crc* in the Δ*lasR*Δ*cbrB*Δ*crc* strain, and pyocyanin levels resembled that of the Δ*lasR*Δ*cbrB* strain with low to undetectable amounts and no significant difference between mono- and co-culture levels (Fig. 4D). Together, these data indicate that relief from catabolite repression promotes interstrain interactions that increase pyocyanin production by theΔ*lasR* strain. We propose the induction of RhlR signalling and its downstream targets are the result of a metabolite present in co-culture that requires CbrB for uptake or catabolism.

### Citrate secreted by wild type can support growth and RhlR induction of LasR- strains

Citrate is a metabolite that requires CbrB for utilization, it is secreted by *P. aeruginosa*, and differentially taken up from rich medium by LasR+ and LasR- strains. *lasR* mutant derivatives lacking either the gene encoding a porin shown to transport citrate (OpdH) or the TctABC citrate transporter did not exhibit significant co-culture phenazine induction (Fig. 4D), and the defect in co-culture phenazine production by the Δ*lasR*Δ*opdH* strain was rescued by *opdH* complementation (Fig. 4D).

To directly explore the possibility that Δ*lasR* mutants were able to use *P. aeruginosa* secreted citrate, we designed an experiment where only the wild type can grow on the provided growth substrate, and Δ*lasR* strain growth can only occur on products released by the wild type. To do this, we used choline as a carbon source (Fig. 5A). Both free choline and choline incorporated into eukaryotic lipids are abundant in infections (44, 45). For these experiments, we engineered a Δ*lasR* strain that lacked *betA* and *betB* which encode a choline oxidase and betaine aldehyde dehydrogenase (46), respectively, that are essential for catabolism of choline as a growth substrate. To test if wild type-secreted products could supportΔ*lasR*Δ*betAB* growth in choline medium, we collected supernatant from wild type cultures grown on 20 mM choline and assessed growth of Δ*lasR*Δ*betAB* in the supernatant diluted 1:1 in fresh medium (Fig. 5A). Supernatants from the wild-culture supported a 4-fold increase in Δ*lasRΔbetAB* density on average over the same strain in fresh medium (Fig. 5B). Furthermore, growth of the Δ*lasR*Δ*betAB* strain was no longer observed if Δ*tctED* (necessary for growth on citrate) were also deleted as we observed a 1.4-fold growth enhancement for the Δ*lasR* Δ*betAB*-*tctED* strain in supernatant relative to fresh medium, which was significantly less than the boost observed for the Δ*lasR*Δ*betAB* strain (Fig. 5B). Supplementing the supernatants with glucose supported greater relative growth than the supernatant alone for both the Δ*lasR*Δ*betAB* and the Δ*lasR*Δ*betAB*Δ*tctED* strains whereas citrate supplementation supported greater relative growth of the Δ*lasR*Δ*betAB* strain compared to supernatant alone, but not for the Δ*lasR*Δ*betAB*Δ*tctED* strain, as expected (Fig. 5B). Because supplementation of glucose or citrate supported additional growth, the supernatants were not inherently restrictive nor limiting aside from carbon. The *tctED*-dependent growth enhancement observed for the Δ*lasR*Δ*betAB* strain in the presence of wild-type-secreted factors is consistent with a model of citrate cross-feeding from wild type to Δ*lasR* cells in co-culture.

**FIG 5.**
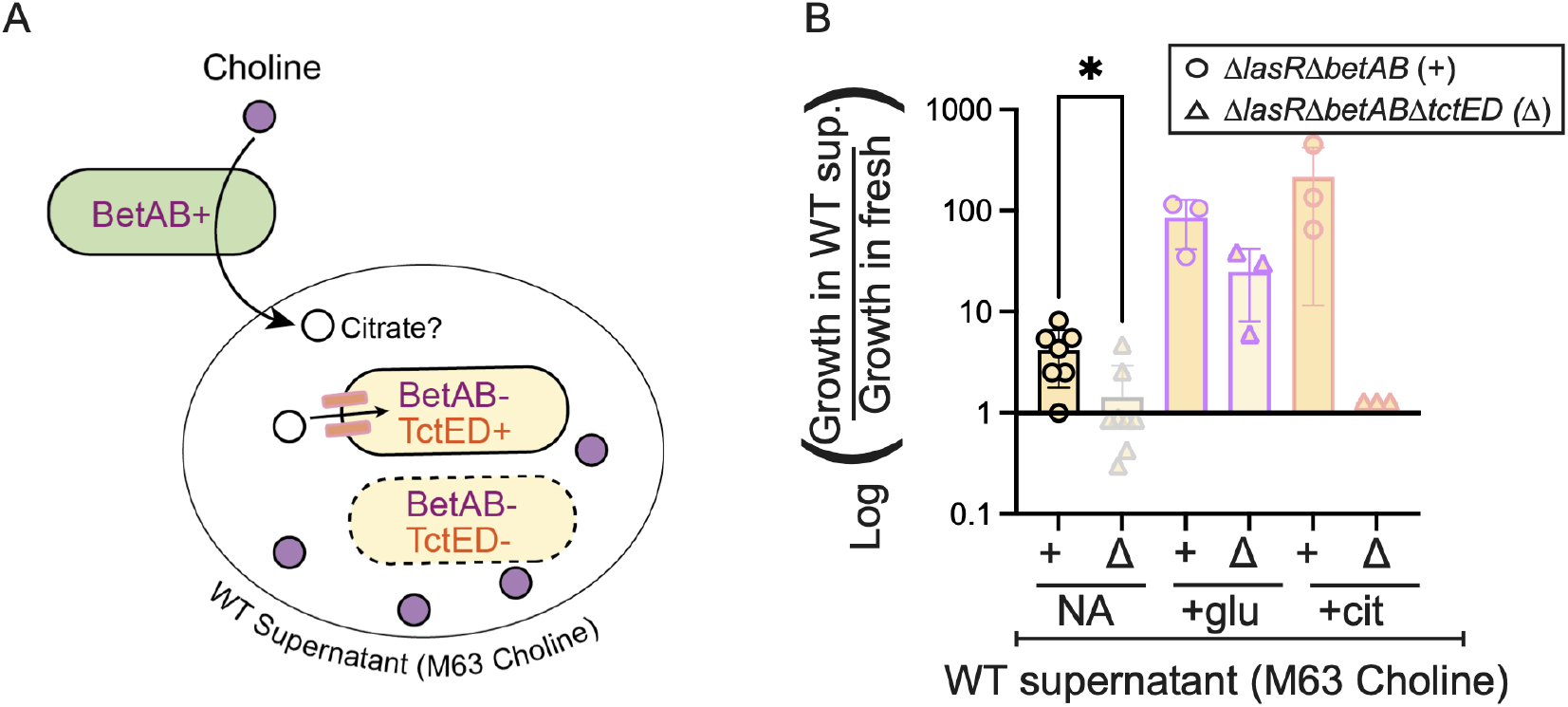
Wild-type secreted factors support *tctED*-dependent growth of *lasR* mutants on an inaccessible carbon source abundant in airway surfactant. A. Schematic of experimetal set up. Choline (purple circle) catabolism requires the enzymes BetA and BetB. Strains lacking these enzymes (BetAB-) do not grow well on choline as a sole carbon source but can consume products released by BetAB+ LasR+ strains (green oval). Citrate (open circle), one secreted metabolite, is taken up by LasR- BetAB-cells (beige oval) in a TctED-dependent manner. B. Growth of the Δ*lasR*Δ*betAB* (circle, +) or Δ*lasR*Δ*betAB*Δ*tctED* (triangles,Δ) strains lacking the citrate responsive two component system controlling citrate uptake strains in supernatant from wild type cultures on M63 with choline as a sole carbon source diluted 1:2 with fresh choline medium without additional carbon supplements (NA) or with additional glucose (“+ glu “) or citrate (“+ cit “) relative to fresh 10 mM choline medium.

### Citrate cross-feeding benefits LasR- strains during intraspecies co-cultures and promotes pyocyanin production in co-culture with *S. aureus*

In light of the findings that (i) secreted citrate is taken up by Δ*lasR* but not wild-type cells via CbrAB and TctED-dependent mechanisms (Fig. 2C), (ii) that in Δ*lasR* cells, citrate induces RhlR activity (Fig. 4B & C), and (iii) published studies indicating the lack of RhlR activity impairs the survival of Δ*lasR* cells due to low levels of RhlR-regulated resistance against QS-regulated exoproducts released in a process termed “policing” (Fig. 6A) (47), we hypothesized that citrate uptake may play a protective role against RhlR-mediated strain-strain inhibition by activating RhlR in LasR- strains—a model we term “protective cross-feeding” (Fig. 6A). Using the experimental system which showed that CbrAB was necessary for the rise of LasR- strains *in vitro* (18), we tested the importance of CbrB-dependent citrate uptake for *lasR* mutant selection. We monitored the rise of LasR- strains in a Δ*tctABC* strain, which showed that LasR-lineages arose more slowly reaching significantly lower proportions of the population when citrate uptake was hindered (Fig. 6B). Consistent with studies describing a RhlR-dependent mechanism to restrict LasR- strain frequency, our previously published data utilizing the same evolution regime depict a more rapid rise of LasR- strains when initiated with a Δ*rhlR* strain (Fig. 6B).

**FIG 6.**
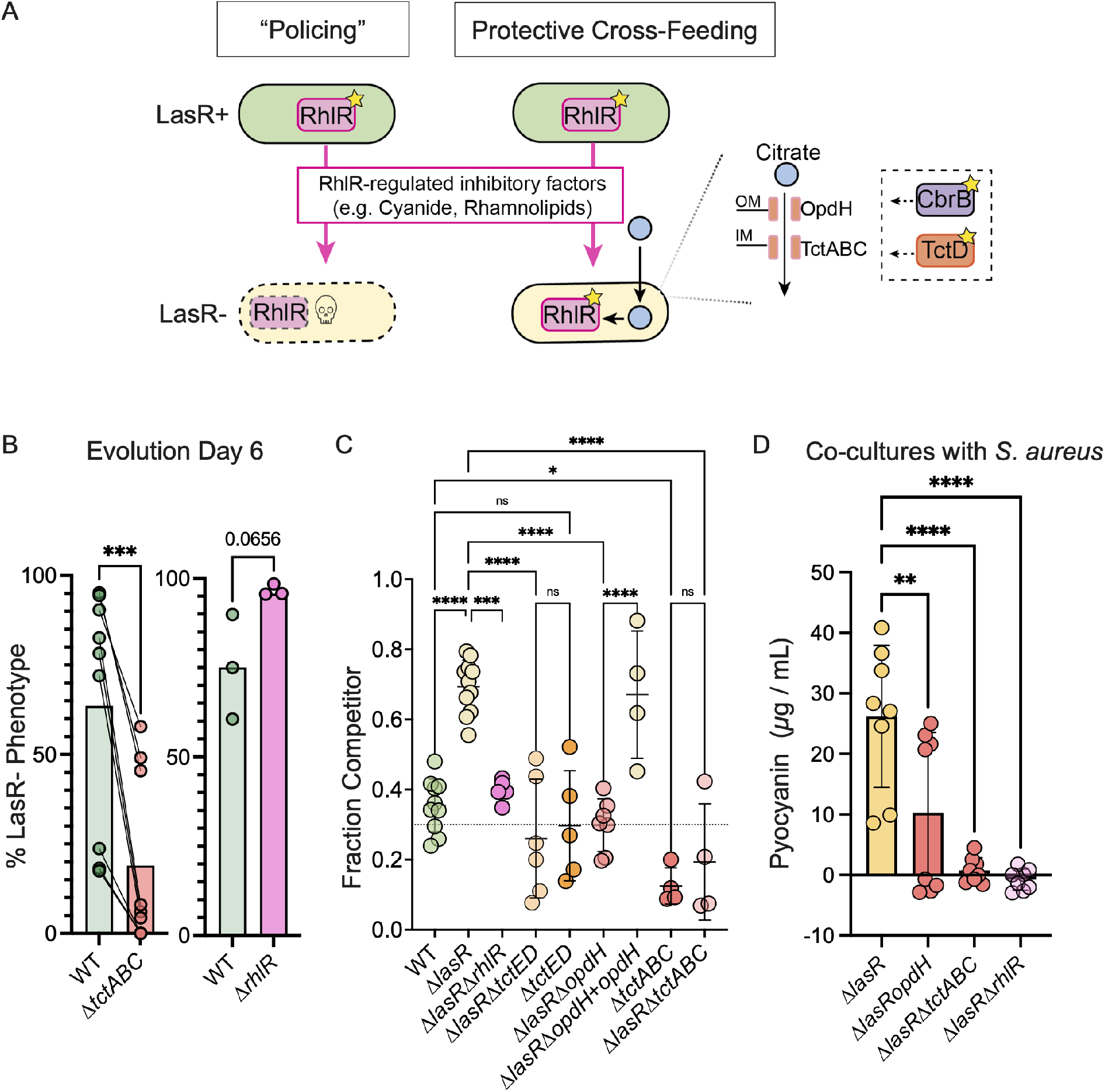
Citrate uptake and RhlR activation impact Δ*lasR* mutant fitness. A. Model illustrating proposed RhlR-dependent interactions in the absence and presence of cross-feeding. Activity of the quorum sensing regulator RhlR (pink rectangle with star indicating activation) reduces sensitivity to RhlR-regulated exoproducts. LasR+ strains (green oval) with active RhlR can “police “, or restrict, the rise of LasR- strains (beige oval) due to differences in RhlR activity. However, citrate cross-feeding from wild type (or other neighboring organisms) and uptake by LasR- strains via the OpdH outermembrane (OM) porin and TctABC innermembrane (IM) transporter under TctD (orange rectangle) transcriptional control and CbrB (purple rectangle) post-transcriptional control promote RhlR-dependent fitness and survival of LasR- strains in a process we term “protective cross-feeding “. B. Percent of population with LasR loss-of-function phenotypes at Day 6 after two passages in LB over an experimental evolution originating from a PA14 wild type,Δ*tctABC*, or *rhlR* ancestor. The data presented for *rhlR* are from our previously published report (18) utilizing the same evolution regime. C. Competitive fitness of Δ*lasR*,Δ*lasR*Δ*rhlR*,Δ*lasR*Δ*tctED*,Δ*tctED*,Δ*lasR*Δ*opdH*, Δ*lasRΔopdH* + *opdH, tctABC, lasR tctABC* competitor strains after 16 h growth on LB agar as colony biofilms when initiated at 30 percent of the starting inoculum (dotted line) with the neutrally tagged wild type (*att*::*lacZ*) making up the difference. D. Pyocyanin levels for the indicatedΔ*lasR* mutants in co-culture with *Staphylococcus aureus*.

Given RhlR has been implicated in the competitive fitness of Δ*lasR* mutant clinical isolates (48, 49), and the presence of LasR+ cells induces RhlR activity in LasR- strains in a TctED-dependent manner (Fig. 4A), we directly assessed the competitive fitness of the Δ*lasR* Δ*rhlR* strain alongside those deficient in citrate uptake relative to the Δ*lasR* strain. Consistent with the observed rise of LasR- strains over the course of serial passaging and previous demonstrations (32, 33), theΔ*lasR* strain was more fit than the wild type in competition assays with a “wild-type” strain harboring a constitutive *lacZ* marker. The Δ*lasR* strain showed a significant increase in competitive fitness with the percentage of the Δ*lasR* competitor rising from 30% of the population at the start of the experiment to 70% after 16 hours (Fig. 6C) while the untagged wild-type control remained close to the starting fraction (0.3) when grown with the tagged strain (Fig. 6C). The Δ*lasRΔrhlR* strain, however, was significantly less fit than theΔ*lasR* strain, with the fraction ofΔ*lasR* Δ*rhlR* cells remaining close to the fraction of the starting inoculum (Fig. 6C). Interestingly, the fitness advantage of theΔ*lasR* mutant was not observed in theΔ*lasR* Δ*tctED*,Δ*lasR* Δ*tctABC*, orΔ*lasR* Δ*opdH* strains with altered citrate consumption (Fig. 6C), and the competitive fitness of the Δ*lasRΔopdH* strain was restored when *opdH* was complemented back onto the chromosome (Δ*lasR* Δ*opdH*+*opdH*) (Fig. 6C). We did not observe any deficit in growth yield or rate between *lasR* mutant derivatives when grown as monocultures to suggest the fitness cost may be specific to co-culture (Fig. S3D). In contrast to the TctED-dependent competitive fitness advantage of LasR- strains, Δ*tctED* in the LasR+ background did not have a fitness cost whereas loss of TctABC did impact LasR+ fitness as well (Fig. 6C). Because RhlR is induced by LasR-cells in co-culture with wild type (Fig. 4A & Fig. S4), and RhlR is important for the competitive fitness of the Δ*lasR* strain, those cells that do not induce RhlR in co-culture may be out-competed.

Previous studies showed that RhlR-regulated factors also contribute to *P. aeruginosa-S. aureus* interactions (as reviewed in (50)). An in-depth analysis of the *P. aeuginosa*-*Staphylococcus aureus* interactions by Zarrella and Khare showed that wild-type *S. aureus* strains had supernatant citrate concentrations in the 100-400 *μ*M range, and that *S. aureus* supernatants induced *opdH* in *P. aeruginosa* (51). Thus, we sought to determine if citrate uptake by Δ*lasR* strains contributed to these interactions as well. *S. aureus* was spread plated on Tryptic Soy Agar (TSA) medium to initiate an even lawn, spot inoculated with *P. aeruginosa* to initiate colony biofilm growth, then co-incubated. We found that while the Δ*lasR* strain showed strong blue-green pigmentation attributable to phenazines, the Δ*lasR*Δ*rhlR*,Δ*lasR*Δ*tctABC*, andΔ*lasR*Δ*opdH* strains did not. Upon quantification, we found significantly less pyocyanin when *S. aureus* was co-cultured with Δ*lasR* mutant derivatives lacking the ability to consume citrate or activate RhlR. Together, these data show that citrate uptake by *P. aerugionsa* contibutes to RhlR-dependent interactions between genotypes of *P. aeruginosa* and across species in *P. aeruginosa-S. aureus* interactions.

## DISCUSSION

In complex environments containing multiple different carbon sources, genetically distinct strains or organisms capable of metabolite secretion, cells can make different choices as to their preferred carbon sources. The differences in metabolic choices between strains can affect important clinical outcomes through virulence factor production and lead to interesting phenomenon such as intraspecies cross-feeding as investigated here between cells with distinct quorum sensing function. Though the mechanism by which *lasR* mutants exhibit elevated CbrB activity remains unclear, the resulting shift toward a more de-repressed metabolism alters how Δ*lasR* cells interact, and we propose this is largely driven by differential metabolite consumption wherein LasR-cells consume citrate, for example, when wild type cells do not (Fig. 1E).

Both wild type and *lasR* mutants are capable of citrate secretion. The loss of *cbrB* in wild type showed no significant change in relative extracellular citrate levels while the same deletion in the Δ*lasR* strain background resulted in more relative extracellular citrate, indicative of differences in consumption rather than secretion (Fig. 2D). The net secretion of citrate by wild type cells could be the result of noisy metabolism inducing metabolite leakage (52), iron limitation promoting release of citrate as a metal scavenging molecule (53), or iron overload as with the IceT transport protein in *Salmonella* wherein citrate is used as a vesicle to secrete excess iron (54). We have previously reported that the siderophore pyochelin promotes citrate release by wild type cells (32) to suggest secretion may relate to the iron content or the signaling capacity of pyochelin itself (55). However, pyochelin supplementation is not necessary for citrate secretion.

Once secreted, citrate becomes an option for neighboring cells where its uptake depends on their metabolic capabilities and preferences. In complex media, LasR- strains consumed more citrate than their LasR+ counterparts (Fig. 1E) and this depended on the CbrAB two component system (Fig. 2C) that enables preferential metabolite consumption. Given the catabolite repression system functions post transcriptionally, differences in activity will result from differences in both transcription and translation.

Citrate can induce a diverse set of transport gene expression—some of which are under the control of the TctED two component system (39, 38, 40). TctED positively regulates the expression of *opdH* and the co-operonic *tctABC* genes encoding an inner-membrane transporter important for growth on citrate (Fig. 3A & Fig. S3ABC) and aconitate (39). Consistent with Underhill et al. (39), the tricarboxylate responsive outer-membrane porin OpdH was not required for growth on citrate (Fig. S3B). Though outer membrane porins are required for citrate to enter the cell, multiple porins are capable of performing this function (56). Once in the periplasm, multiple tricarboxylate transporters encoded with the genome can bring tricarboxylates across the inner-membrane (39). While no significant differences were observed in TctD transcriptional activity between wild type and the Δ*lasR* strain, *opdH* promoter activity in theΔ*lasR* strain trended lower which may relate to the increased consumption by LasR- strains and implicate extracellular or periplasmic citrate levels, rather than intracellular, as an inducer of TctED activity. In co-culture with wild-type cells which secrete upwards of 300 μM citrate, we observed small but significant increases in TctD-dependent *opdH* promoter activity in co-culture (Fig. 3C). In a prior assessment of all TCA intermediates, *opdH* expression was most strongly stimulated by growth on citrate (38). Once transcribed, these gene products are likely susceptible to translational repression via the catabolite repression system for reasons previously described.

When activated by RhlI-synthesized C4HSL, RhlR is important for competitive fitness (47, 57, 48) in strain PAO1 among other strains. Here, we extend results to show the requirement of RhlR in the PA14 Δ*lasR* when co-cultured with the wild type (Fig. 6C). Citrate induction of RhlI-RhlR signaling may increase *lasR* mutant fitness in LasR+/LasR-co-cultures by increasing its resistance to RhlR-controlled exoproducts (e.g. hydrogen cyanide, rhamnolipids, or other secreted factors (47, 58)) (Fig. 6A). The induction of *rhlI* promoter activity required tricarboxylate transport machinery (Fig. 4C & Fig. S3C), and the differential citrate uptake likely explains why citrate induces activity of RhlR in LasR-but not LasR+ strains (32). Consistent with this model, we observed a fitness deficit in Δ*lasR* mutants lacking tricarboxylate regulatory components, uptake machinery or RhlR itself when grown in the presence of wild type (Fig. 6C), but not in isolation (Fig. S3D). However, it is possible that citrate uptake is important for other reasons (e.g. metal uptake), and that other metabolites may participate in the activation of RhlR in LasR- strains.

The intracellular metabolomes of LasR-cells are distinct from LasR+ cells regardless of strain background and medium type (Fig. 1A). The higher intracellular citrate observed in LasR- strains in this study parallels metabolomics analysis of a strain (Δ*lasI*Δ*rhlI*) lacking the ability to produce the QS signals, referred to as autoinducers, suggesting differences in citrate are linked to quorum sensing activity more broadly (17). The distinct metabolic states induced by quorum sensing activity highlight the potential for similar cross-feeding interactions to occur within genetically identical or clonal populations where distinct clusters of cells with differential quorum sensing activity exist. Heterogeneity in quorum sensing signalling has been well documented (59, 60, 61), lending support to this possibility, and extending the implications beyond genetically distinct cells to identical ones with distinct expression profiles. Beyond *P. aeruginosa*, many other cell types (host and microbial) secrete citrate. Thus, the cross-feeding in-teractions described here likely extend to those between other microbes and host cells where virulence factor production is differentially stimulated between LasR- and LasR+ strains (22, 24, 23). We observed a TctED-dependent phenazine interaction between *S. aureus* and *lasR* loss-of-function mutants (Fig. 6D). Furthermore, like citrate, many of the other metabolites commonly secreted as a result of overflow metabolism or the Warburg effect (62, 37, 63), including pyruvate and acetate, require CbrB for consumption and/or metabolism (64). Metabolite exchange can allow new microbial inhabitants to thrive, alter ecosystem structure and influence other emergent properties or functions specific to high density cells. Additional work will be required to elucidate how the intraspecies interactions described here may shape community structure in evolving *P. aeruginosa* populations and its interactions with neighboring cells.

## MATERIALS AND METHODS

### Strains and Growth Conditions

Strains used in this study are listed in Table S2. *P. aeruginosa* strains were maintained on LB (lysogeny broth) with 1.5% agar. *S. aureus* was maintained in TSB (Tryptic Soy Broth) with 1.5% agar. Yeast strains for cloning were maintained on YPD (yeast extract, peptone, dextrose per L) with 2% agar at 30 °C. Planktonic bacterial cultures (5 mL) were grown on a roller drum at 37°C in 13 mm borosilicate tubes at 37 °C.

### Strain Construction

Plasmids were constructed using a *Saccharomyces cerevisiae* recombination technique described previously (65). The construct for constitutive mKate2 fluorescence was constructed as described in Kassety et al. with two tandem codon optimized mKate2 under a synthetic tac promoter (66). Plasmid candidates were selected by restriction digest, and successful candidate plasmids were sequenced for verification of the insert. In frame-deletions, complementation constructs, and integrated promoter fusions were introduced into *P. aeruginosa* by conjugation via S17/lambda pir *E. coli*. Merodiploids were selected by drug resistance, and double recombinants were obtained by sucrose counter-selection and genotype screening by PCR.

### Metabolomics

Strains were grown overnight in 5 mL of LB broth for 16 h from single colonies taken from freshly struck strains on LB plates, and 5 *μ*L of the overnight culture was used to initiate colony biofilm growth on LB agar or artificial sputum medium (ASM) agar made as described in (33). For each replicate, four plates with 18 colony biofilms each were pooled during collection after 16 h of growth at 37 °C. Colonies were scraped from the agar surface using a rubber policeman, deposited in a 1.5 mL tube, briefly pelleted, stored at −80 °C, and sent to Metabolon (Morrisville, NC) for metabolomics analysis. Five replicates were collected for each sample/condition. Raw counts were scaled and imputed, then normalized using the total raw counts across all metabolites by sample. Pareto normalized metabolite counts were used as input for principal component analysis using the R package ggfortify. The ggplot2 and ggprism R packages were used to visualize the differential metabolite data using R (Version 4.2.1) (67, 68). Only metabolites detected across all samples in comparison were included in volcano plots.

### Citrate Quantification

For extracellular citrate measurements, 5 mL of LB was incoculated with a single colony, grown for 16 h, pelleted by 10 min at max speed, and filtered through a 0.22 *μ*m pore-size polycarbonate filter. Citrate was measured as previously (32) using½ reactions in a commercially available Megazyme citric acid enzymatic kit (cat. K-CITR). A citrate standard was included in every assay. The citrate concentration is reported relative to the cellular density (OD_600 *nm*_) measured after 16 h of growth at 37 °C on a roller drum where indicated. For citrate consumption assays, a 16 h LB culture was adjusted to an OD_600 *nm*_ of 1.5 in 2 mL total volume of LB and incubated at 37 °C for 10 min after which 2 mM citrate was added. Cultures were incubated statically at 37 °C for 24 h with Parafilm covering the uncapped tube opening to allow gas exchange and reduce evaporation. After incubation cells were pelleted and filtered for assessment in the enzymatic assay as described above.

### Growth Assays

Growth was assessed in M63 base (10 g (NH_4_)_2_SO_4_, 15 g KH_2_PO_4_, 35 g K_2_HPO_4_ per liter) containing 10 mM citrate in two distinct ways. Data shown in Fig. 2B was collected using the method described in (18). In brief, 5 mL LB cultures grown for 16h were adjusted to an OD_600 *nm*_ of 1 in LB and 250 *μ*L of the normalized culture was added to 5 mL of fresh M63 base containing 10 mM disodium citrate (ACROS) as a sole carbon source. The OD_600 *nm*_ was monitored using a Spectronic 20D+ (Spec20) during growth on a roller drum at 37 °C. Growth reported in Fig. S3B were performed by inoculating single colonies from freshly struck LB plates into 5 mL of M63 with 10 mM disodium citrate (ACROS). The density (OD_600 *nm*_) was measured in a 1 cm cuvette with a 1:10 dilution after 24 h of incubation on a roller drum at 37 °C.

### Beta-galactosidase (*β*-gal) Quantification

Cells with a promoter fusion to *lacZ*-GFP integrated at the *att* locus were grown in 5 mL cultures of LB at 37 °C for 16 h. The cultures were diluted to a starting OD_600 *nm*_ of 1 in LB. For assessments as colony biofilms (as in Fig. 3A & B and Fig. 4B & C), a 5 *μ*L aliquot of each normalized culture was spotted onto LB agar plates with the indicated concentration of disodium citrate (pH 7). After 24 h at 37 °C, the colony biofilms were cored, re-suspended in 500 *μ*L of LB by vigorous shaking on a Genie Disrupter for 5 min as previously described (32), and the cell suspension was used as input into the *β*-Gal assay. *β* Gal activity was measured as described by Miller using the equation: Miller Unit= 1000 ((*OD*_420*nm*_ − (1.75 *OD*_550*nm*_))/(*t ime volume OD*_600*nm*_), where time is reported in minutes and volume in milliliters (69). For co-culture induction assays, a 250 *μ*L aliquot of normalized culture was added to 5 mL of fresh LB to initiate monocultures and incubated on a roller drum for 6 h at 37 °C. For co-cultures, the inoculation was initiated in a 8:2 ratio of the PA14 *att*:P*tac*-mKate strain to the Δ*lasR* derivative containing the designated promoter fusion. To do this, we added 200 *μ*L of the PA14 *att:*P*tac*-mKate strain and 50 *μ*L of the *lasR* mutant to 5 mL of LB for comparison with the monocultures where 250 *μ*L of OD_600 *nm*_ = 1 normalized cells were added. After 6 h, we collected the cultures and adjusted the optical density to 0.5 for use in the *β*-Gal assay. An appropriate dilution was plated for each co-culture onto LB plates containing 150 *μ*g / mL 5-bromo-4-chloro-3-indolyl-D-galactopyranoside (X-Gal) using glass beads, and the plates were incubated at 37 ° C until blue colonies were visible ( ~24 h). To calculate the relative *β*-Gal activity, the OD_600 *nm*_ value in the Miller unit equation was multiplied by the fraction containing the *lacZ* marker as indicated by blue:white colony counts. Each experiment was repeated in at least four independent experiments.

### Pyocyanin Quantification

Overnight cultures were normalized in LB to an OD_600 *nm*_ = 1. For monocultures, a 3 *μ*L aliquot of OD-adjusted cultures was plated in four replicate wells of a 96-well plate in which each well was filled with 200 *μ*L of LB agar. For co-cultures, 700 *μ*L of the Δ*phz* strain was mixed with 300 *μ*L of the indicated Δ*lasR* mutant, vortexed, and then a 3 *μ*L aliquot of the mixed culture was plated in four replicate wells of the same 96-well plate as the monocultures. After 16 h of incubation at 37 °C, the 96-well plates were placed at −80 °C until extraction. For pyocyanin extraction, two agar plugs were placed in a 2 mL eppendorf tube with 500 *μ*L of chloroform and agitated on a Genie Disruptor (Zymo) for 2 min. After which 200 *μ*L of the bottom chloroform layer was placed into a fresh 1.5 mL tube, and the chloroform extraction was repeated with 500 *μ*L of fresh chloroform. Another 300 *μ*L of the bottom layer was added to the previous 200 *μ*L aliquot. To the chloroform layer, 500 *μ*L of 0.2 N HCl was added to acidify the sample turning the pyocyanin from blue to pink, and the optical density was measured at 520 nm alongside an acidified pyocyanin standard to determine the concentration in *μ*g / mL for each strain by interpolation. The pyocyanin extracted was reported per colony biofilm (or agar plug).

### Flow Cytometry

Strains with the promoter from the RhlR-regulated gene *rhlI* fused to green fluorescent protein (GFP) and the *lacZ* gene were grown in monoculture or in co-culture with a “wild type” strain constitutively expressing the mKate2 fluorophore under a synthetic *tac* promoter, referred to as the PA14 att:P*tac*-mKate strain. Overnight LB cultures of the PA14 att:P*tac*-mKate strain and the designated *lasR* mutants with the *lacZ* – GFP fusion integrated at the Tn7 *att* site were adjusted to an OD_600 *nm*_ of 1. For monocultures, a 250 *μ*L aliquot was added to 5 mL fresh LB. For co-cultures, 200 *μ*L of the PA14 att:Ptac-mKate and 50 *μ*L of the designated *lasR* mutant were added to 5 mL of fresh LB broth. After 6 h of incubation on a roller drum at 37 °C, a 500 *μ*L aliquot of subculture was pelleted for 5 min at 13,000 RPM, resuspended in 500 *μ*L of PBS + 0.01% TritonX100, diluted 100 fold in PBS + 0.01% TritonX100, and diluted again 1:1 in PBS without detergent. The diluted cells were placed on ice until processed by flow cytometry. The data were collected by Beckman Coulter Cytoflex S and analyzed with FlowJo version 10.8.1. In short, single cell gating was done using FSC versus SSC and cells without mKate expression were gated using the ECD channel. GFP expression was quantified by FITC Median Fluorescence Intensity of the single cell, mKate negative populations.

### Syntrophic Growth

Single colonies of PA14 wild type were inoculated into a 250 mL baffled flask containing 100 mL of M63 base with 20 mM choline chloride. After 48 h on a shaker (240 RPM) at 37 °C, cultures were pelleted at max speed for 20 minutes in 50 mL falcon tubes. The supernatant was filtered through a 0.22 *μ*m pore size polycarbonate filter and stored at 4 °C. Supernatant was diluted 1:1 in fresh choline medium with 2 mL per well in a 12-well plate. Colonies of the Δ*lasR*Δ*betAB* orΔ*lasR*Δ*betAB*Δ*tctED* strain from a freshly struck LB plate were re-suspended in water and adjusted to an OD_600 *nm*_ of 2. The cell re-suspension was used to inoculate the diluted supernatant or fresh choline medium at a starting OD_600 *nm*_ ∼ 0.05, and growth monitored after 48 h by CFU count in n > 3 experiments and/or OD_600 *nm*_ in two experiments. Supernatant from experiments performed on at least three distinct days were tested for growth enhancement in at least three separate experiments.

### Competition Assays

Competition assays were performed as in (32). In summary, strains were grown for 16 h from a single colony in a 5 mL LB culture on a roller drum at 37 °C. Based on a 1:10 dilution of the 16 h culture in a 1 cm cuvette, cultures were adjusted to OD_600 *nm*_ = 1 in the required volume of LB. Density adjusted competitor strains were mixed with PA14 *att*::*lacZ* strain in a 7:3 ratio by adding 700 *μ*L of the PA14*att*::*lacZ* strain with 300 *μ*L of the competitor strain. Following a brief vortex, a 5 *μ*L aliquot of the combined suspension was spotted on LB agar. After 16 h, colony biofilms (and agar) were cored using the back of a sterile P1000 pipette tip, placed in 1.5 mL tubes with 500 *μ*L LB, and shaken vigorously for 5 min using a Genie Disruptor (Zymo). This suspension was diluted in LB, spread on LB plates supplemented with 150 *μ*g / mL 5-bromo-4-chloro-3-indolyl-D-galactopyranoside (X-Gal) using glass beads, and incubated at 37 °C until blue colonies were visible ( ~24 h). The number of blue and white colonies per plate were counted and the final proportions recorded. Each competition was replicated on 4 - 7 separate days.

### *Staphylococcus aureus* Interactions Assay

Protocol was based off of (25) with modifications. *S. aureus* strain SH1000 was grown in 5 mL of TSB at 37 °C for 16 h and adjusted to OD_600 *nm*_ = 0.1 in the required volume of TSB. A ∼ 100 or 150 uL aliquot of density-adjusted *S. aureus* culture was spread on a Tryptic Soy Agar (TSA) plate with sterile glass beads and allowed to dry in a biosafety cabinet for ~10 min. *P. aeruginosa* cultures (16 h, 5 mL LB) were adjusted to OD_600 *nm*_ = 1 in LB, and a 5 *μ*L aliquot was spotted onto the the lawn of *S. aureus* after the drying period to prevent colony spreading. After 16 h incubation at 37 °C, plates were placed at room temperature and imaged daily with a Canon EOS Rebel T6i digital camera. The phenazine pyocyanin was extracted from cored *P. aeruginosa* colonies and the underlying *S. aureus* lawn on day 3 and quantified as described previously. Each experiment was repeated on at least three independent days.

### Experimental Evolution

Experimental evolution was performed exactly as described in (18). In brief, an LB culture was inoculated with a single colony of the indicated strain, grown for 24 h, and normalized to an OD_600 *nm*_ of 1 in LB. This normalized culture was used to begin three independent lines with a starting OD_600 *nm*_ of 0.05. These three lines were passaged every 48 h with 25 *μ*l being transferred to 5 ml of fresh LB at each passage. At each time point, a 35 - 50 *μ*L aliquot at a 10 5 dilution was plated onto LB agar and the percent of colonies with a sheen LasR-phenotype due to HHQ accumulation were enumerated. Each evolution was repeated three times (with three lines each) and counted by an independent enumerator.

### Availability of data and materials

All data from metabolomics analysis, including raw and normalized counts, are included in Table S1. All materials and data are included in the manuscript or supplemental material, or will be made available in a timely fashion, at reasonable cost, and in limited quantities to members of the scientific community for noncommercial purposes, subject to requirements or limitations imposed by local and/or U.S. Government laws and regulations.

## SUPPLEMENTAL MATERIAL

The following supplemental material are included:

**TABLE S1**. Relative intracellular metabolites detected for LasR+ and LasR-paired isolates on LB or ASM.

**FIG S1**. Citrate is the only tricarboxylic acid (TCA) cycle intermediate significantly enriched in LasR-cells across genetic background and media type. Volcano plots showing differential intracellular metabolite counts (log_2_) for the laboratory isolates PA14 and Δ*lasR* on Artificial Sputum Medium (ASM) (A) or the LasR-clinical isolate compared to its closely related LasR+ progenitor on LB (B) relative to the log_10_ (P value). Metabolites are colored based on their categorization as dipeptides (lavender) or inclusion in the TCA cycle (light blue). 4-hydroxy-2-heptylquinoline (HHQ) and N-butanoyl-L-homoserine lactone (C4HSL) which are known to be enriched and depleted in Δ*lasR* mutants, respectively are also indicated. C4HSL was not significantly changed in the ASM dataset between strains. C. The log_2_ fold change and statistical significance of metabolites noted for LasR-over LasR+ strains in all three conditions tested (3 circles, left to right): PA14Δ*lasR* / WT on LB, LasR-DH2415/ LasR+ DH2417 on LB, and Δ*lasR* / WT on artificial sputum medium (ASM). Circle fill is colored by relative log_2_ fold change as indicated by scale ranging from −2 to 2. Significance as determined by Welch’s t-test is indicated by circle outline (P value ≤ 0.05, solid outline and P value > 0.05 grey, dotted). Metabolites listed in grey text without numbers were not detected in our analysis.

**FIG S2**. Total extracellular citrate in cell-free supernatant taken from 5 mL LB cultures of PA14 wild type or the Δ*lasR* strain compared to the uninoculated LB medium blank. Statistical significance as determined by One-Way ANOVA with Dunnett’s mul-tiple comparison test, ****, P value < 0.0001 and ns, not significant (P value = 0.9902 for the LB andΔ*lasR* comparison).

**FIG S3**. TctED positively regulate citrate transport necessary for growth on citrate but not LB. A. Model illustrating the positive control both the sensor kinase TctE and response regulator TctD have on select genes involved in tricarboxylate transport (i.e. *opdH* and *tctABC*) in response to citrate. OpdH is a porin located in the outermembrane and TctABC is an innermembrane (IM) transporter. B. Growth in M63 base containing 10 mM citrate as a sole carbon source after 24 h from a single colony inoculum for the indicated strains. C. Quantification of P*opdH* – GFP – *lacZ* via beta-galactosidase assay for the indicated strains when grown for 24 h as colony biofilms on LB agar or LB agar supplemented with 20 mM citrate. D. Growth yield after 8 h subculture and exponential growth rate in 5 mL LB cultures.

**FIG S4**. In co-culture with wild type,Δ*lasR* cells exhibit increased RhlR-dependent *rhlI* promoter activity at the cellular level. A. Histogram of mKate2 negative counts by expression of green fluorescent protein detected by FITC of theΔ*lasR* P*rhlI* – GFP – *lacZ* cells in monoculture (light beige, solid line) or co-culture (darker beige, dotted line) with a wildtype strain expressing mKate2 constitutively under the P*tac* promoter at a neutral site (illustrated in inset). Cell counts and median fluorescence intensity (MFI) indicated for cells lacking mKate expression (mKate*OF F*) that exhibit GFP fluorescence (GFP*ON*) after 6h subculture in LB. B. MFI of P*rhlI*– GFP – *lacZ* or the indicated strains when grown alone or in co-culture with the mKate2-tagged wild type after 6 h in LB as described in A. In co-culture with wild type,Δ*lasR* cells exhibit increased *rhlI* promoter activity at the cellular level.

**TABLE S2**. Strains used in this study.

## ACKNOWLEDGMENTS

Research reported in this publication was supported by grants from the Cystic Fibrosis Foundation HOGAN19G0 and NIH/NIAID T32AI007519 (D.L.M). Additional support came from NIGMS P20GM113132 through the Molecular Interactions and Imaging Core (MIIC), STANTO19R0 from the Cystic Fibrosis Foundation and NIDDK P30-DK117469 (Dartmouth Cystic Fibrosis Research Center). We would like to acknowledge Matthew T. Cabeen and lab for sharing strains invaluable for focusing our efforts amidst a redundant transport system as well as Brennan O ‘Toole and Marina Ruzic whose diligent counting efforts added rigor as independent enumerators for LasR-colony phenotypes in the experimental evolution.

