## Supplemental figures for "Citrate cross-feeding between *Pseudomonas aerguinosa* genotypes supports *lasR* mutant fitness"

Mould 2023 Preprint Supplement

**Compiled** May 29, 2023
This is a draft manuscript, pre-submission

### SUPPLEMENTAL FIGURES

The following supplemental materials are associated with the manuscript:

**Table S1** Relative intracellular metabolites detected for LasR+ and LasR- paired isolates on LB or Artificial Sputum Medium (ASM).[a]

[a]See attached excel file with raw and normalized metabolite counts with differential abundance and significance indicated.

Mould 2023 Preprint Supplement Mould 2023 Preprint Supplement Mould 2023 Preprint Supplement Mould 2023 Preprint Supplement Mould 2023 Preprint Supplement



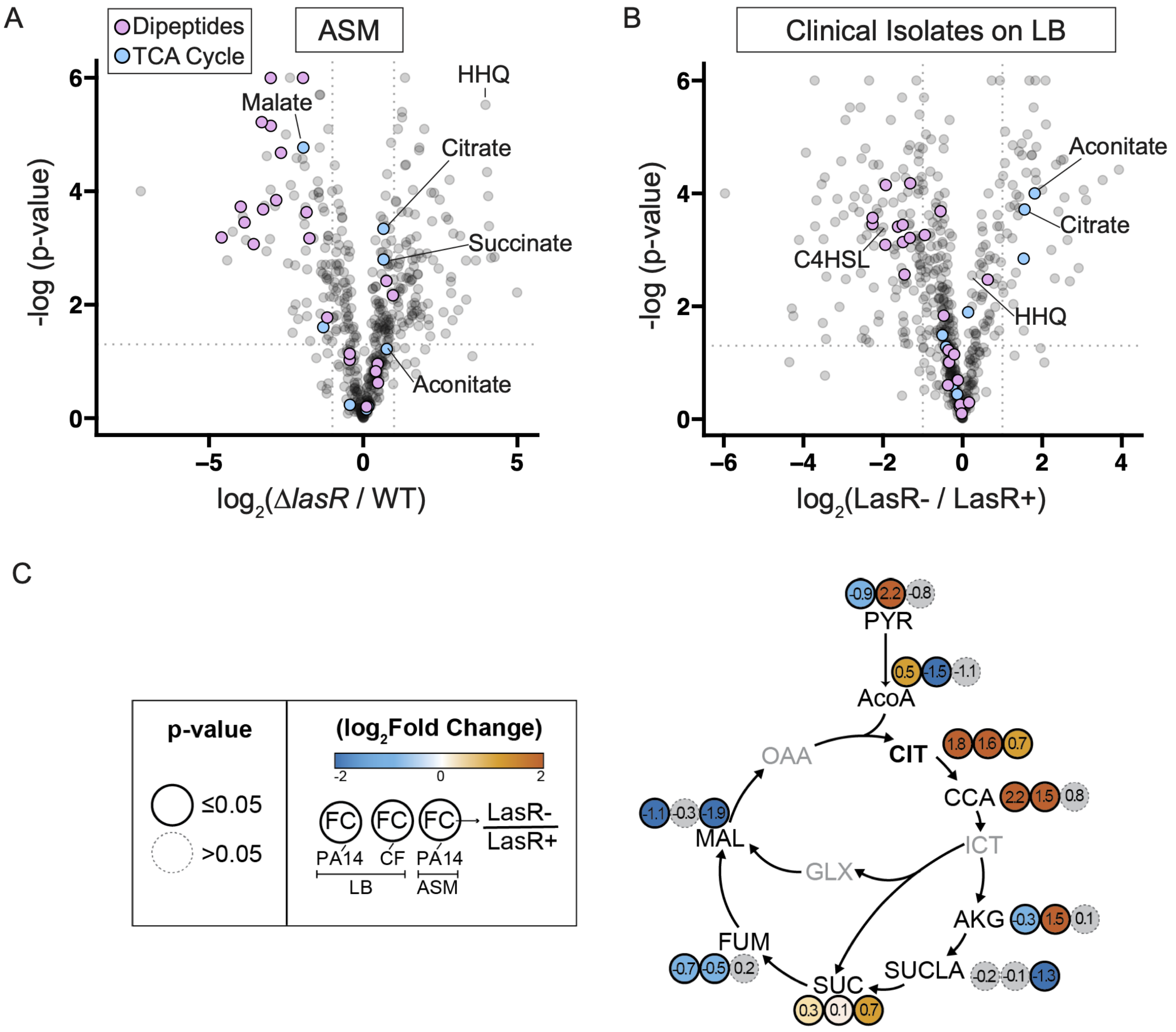


**Fig. S1 Citrate is the only tricarboxylic acid (TCA) cycle intermediate significantly enriched in LasR- cells across genetic background and media type.** Volcano plots showing differential intracellular metabolite counts ($\log_2$) for the laboratory isolates PA14 and Δ*lasR* on Artificial Sputum Medium (ASM) (A) or the LasR- clinical isolate compared to its closely related LasR+ progenitor on LB (B) relative to the $-\log_{10}$ (P value). Metabolites are colored based on their categorization as dipeptides (lavender) or inclusion in the TCA cycle (light blue). 4-hydroxy-2-heptylquinoline (HHQ) and N-butanoyl-L-homoserine lactone (C4HSL) which are known to be enriched and depleted in *lasR* mutants, respectively are also indicated. C4HSL was not significantly changed in the ASM dataset between strains. C. The $\log_2$ fold change and statistical significance of metabolites noted for LasR- over LasR+ strains in all three conditions tested (3 circles, left to right): PA14 Δ*lasR* / WT on LB, LasR- DH2415/ LasR+ DH2417 on LB, and Δ*lasR* / WT on artificial sputum medium (ASM). Circle fill is colored by relative $\log_2$ fold change as indicated by scale ranging from -2 to 2. Significance as determined by Welch's t-test is indicated by circle outline (P value ≤ 0.05, solid outline and P value > 0.05 grey, dotted). Metabolites listed in grey text without numbers were not detected in our analysis.

Mould 2023 Preprint Supplement



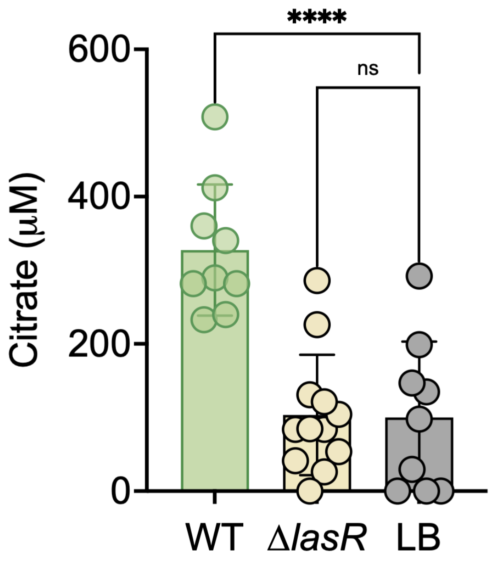


**Fig. S2 Total extracellular citrate in cell-free supernatant taken from 5 mL LB cultures of PA14 wild type or the Δ*lasR* strain compared to the uninoculated LB medium blank.** Statistical significance as determined by One-Way ANOVA with Dunnett's multiple comparison test, ****, P value < 0.0001 and ns, not significant (P value = 0.9902 for the LB and Δ*lasR* comparison).

Mould 2023 Preprint Supplement Mould 2023 Preprint Supplement Mould 2023 Preprint Supplement Mould 2023 Preprint Supplement Mould 2023 Preprint Supplement Mould 2023 Preprint Supplement



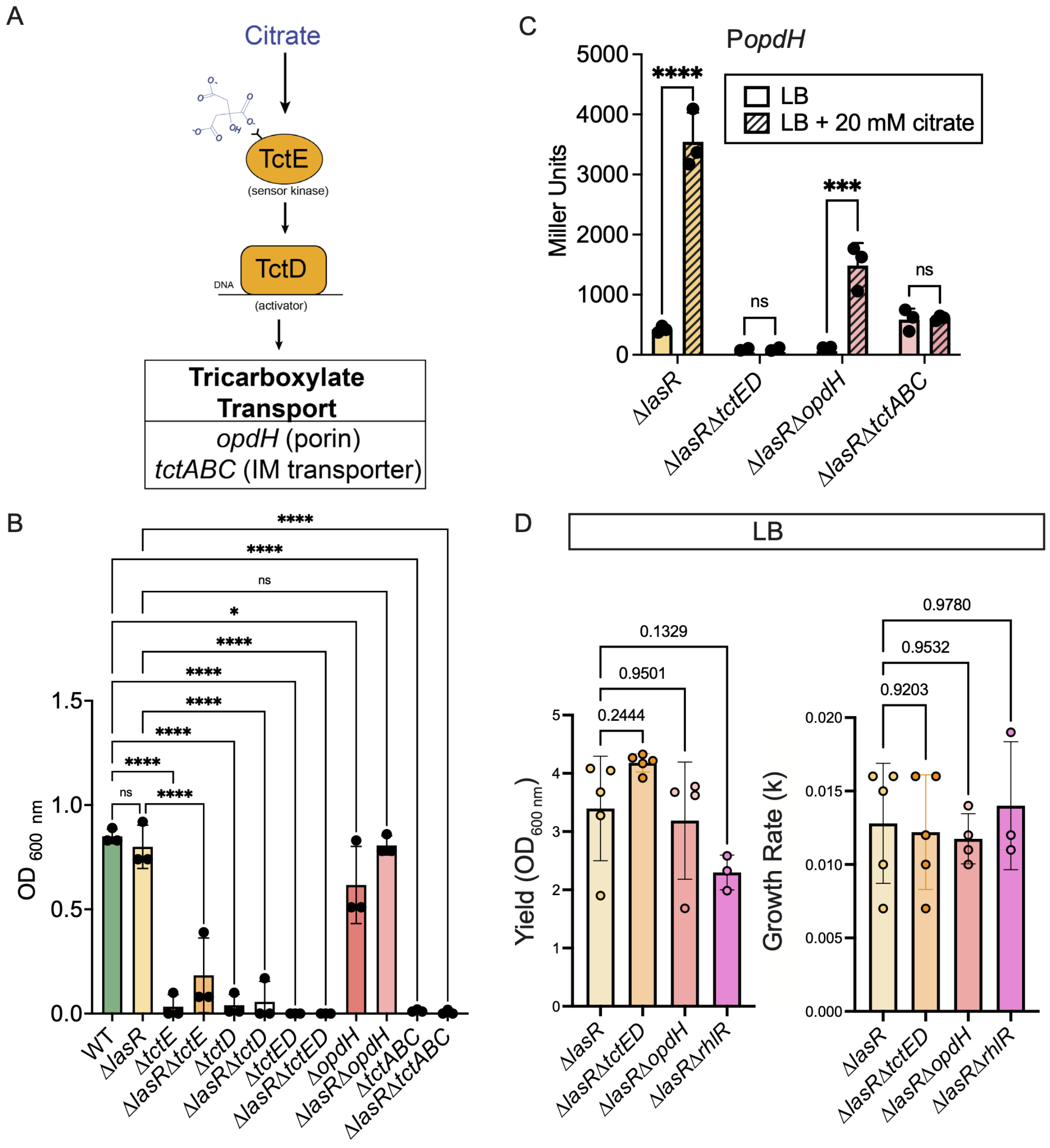


**Fig. S3 TctED positively regulate citrate transport necessary for growth on citrate but not LB.** A. Model illustrating the positive control both the sensor kinase TctE and response regulator TctD have on select genes involved in tricarboxylate transport (i.e. *opdH* and *tctABC*) in response to citrate. OpdH is a porin located in the outermembrane and TctABC is an innermembrane (IM) transporter. B. Growth in M63 base containing 10 mM citrate as a sole carbon source after 24 h from a single colony inoculum for the indicated strains. C. Quantification of P*opdH* – GFP – *lacZ* via beta-galactosidase assay for the indicated strains when grown for 24 h as colony biofilms on LB agar or LB agar supplemented with 20 mM citrate. D. Growth yield after 8 h subculture and exponential growth rate in 5 mL LB cultures.

Mould 2023 Preprint Supplement Mould 2023 Preprint Supplement Mould 2023 Preprint Supplement Mould 2023 Preprint Supplement Mould 2023 Preprint Supplement Mould 2023 Preprint Supplement



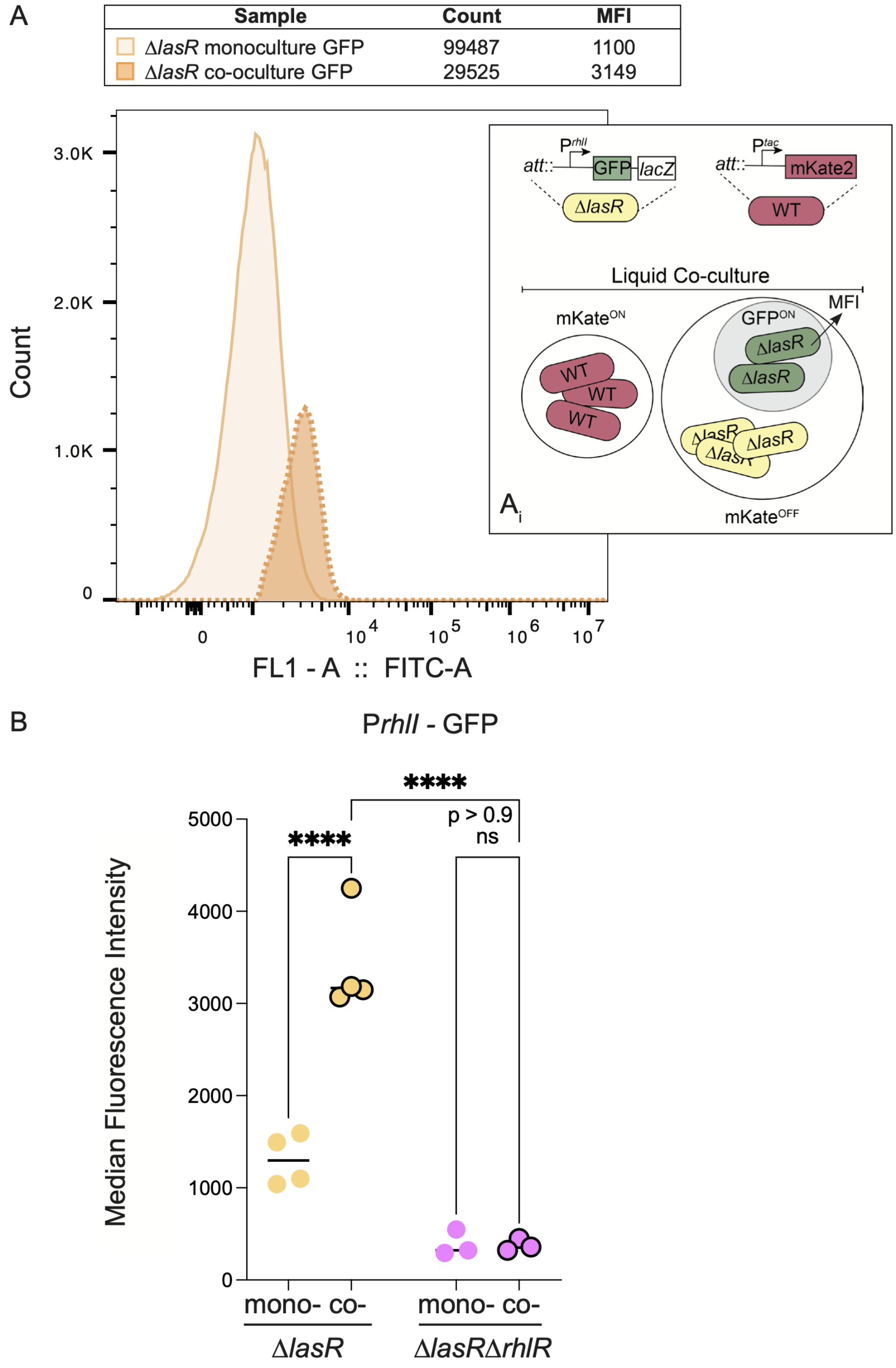


**Fig. S4 In co-culture with wild type, Δ*lasR* cells exhibit increased RhlR-dependent *rhlI* promoter activity at the cellular level.** A. Histogram of mKate2 negative counts by expression of green fluorescent protein detected by FITC of the Δ*lasR* P*rhlI* – GFP – *lacZ* cells in monoculture (light beige, solid line) or co-culture (darker beige, dotted line) with a wildtype strain expressing mKate2 constitutively under the P*tac* promoter at a neutral site (illustrated in inset). Cell counts and median fluorescence intensity (MFI) indicated for cells lacking mKate expression (mKate$^{OFF}$) that exhibit GFP fluorescence (GFP$^{ON}$) after 6h subculture in LB. B. MFI of P*rhlI*– GFP – *lacZ* or the indicated strains when grown alone or in co-culture with the mKate2-tagged wild type after 6 h in LB as described in A. In co-culture with wild type, Δ*lasR* cells exhibit increased *rhlI* promoter activity at the cellular level.
