## Supplementary material for "Citrate cross-feeding between *Pseudomonas aerguinosa* genotypes supports *lasR* mutant fitness": Table S2

**Supplemental Table 2**

**Table S2. Strains and plasmids used in this study**

| *Strain* | *Strain.ID* | *Description* | *Source* |
|---|---|---|---|
| *P. aeruginosa* | | | |
| PA14 WT | DH122 | Laboratory reference strain | (1) |
| PA14 Δ*lasR* | DH164 | DH122 with in-frame deletion of *lasR* (*PA14_45960*) | (2) |
| NC-AMT0101-1-2 | DH2417 | Chronic CF lung infection isolate with functional LasR allele, parent of NC-AMT0101-1 | (3) |
| NC-AMT0101-1-1 | DH2415 | Chronic CF lung infection isolate related to DH2417 with LasR LOF (frameshift) allele | (3) |
| PA14 Δ*tctE* | | PA14 WT (DH122) with in-frame deletion of *tctE* | This study |
| PA14 Δ*lasR*Δ*tctE* | | PA14 Δ*lasR (*DH164) with in-frame deletion of *tctE* | This study |
| PA14 Δ*lasR*Δ*cbrB* | DH3924 | PA14 Δ*lasR* (DH164) with in-frame deletion of *cbrB* (PA14_62540) | (4) |
| PA14 Δ*lasR*Δ*cbrB* + *cbrB* | DH3925 | PA14 Δ*lasR*Δ*cbrB* (DH3924) with complementation of *cbrB* (PA14_62540) at the native locus | (4) |
| PA14 Δ*lasR*Δ*cbrB*Δ*crc* | DH3926 | PA14 Δ*lasR*Δ*cbrB* (DH3924) with in-frame deletion of *crc* (PA14_70390) | (4) |
| PA14 Δ*lasR*Δ*cbrB*Δ*crc* + *crc* | DH3926 | PA14 Δ*lasR*Δ*cbrB*Δ*crc* (DH3926) with complementation of *crc* (PA14_70390) at the native locus | This study |

| | | | |
|---|---|---|---|
| PA14 Δ*lasR* P*rhlI*-GFP-*lacZ* | DH3313 | PA14 Δ*lasR* (DH164) expressing P*rhlI*-GFP-*lacZ* promoter fusion at the *att*::Tn7 site | (5) |
| PA14 Δ*lasR*Δ*cbrB* P*rhlI*-GFP - *lacZ* | | PA14 Δ*lasR*Δ*cbrB* (DH3924) expressing P*rhlI*-GFP-*lacZ* promoter fusion at the *att*::Tn7 site | This study |
| PA14 Δ*phz* | DH933 | In-frame deletions of *phzA1-G1* and *phzA2-G2* | (6) |
| PA14 Δ*lasR*Δ*rhlR* | DH2944 | In-frame deletion of *lasR* and *rhlR* | (7) |
| PA14 WT P*opdH*-GFP-*lacZ* | | PA14 WT (DH122) expressing P*opdHI*-GFP-*lacZ* promoter fusion at the *att*::Tn7 site | This study |
| PA14 Δ*lasR* P*opdH*-GFP-*lacZ* | | PA14 Δ*lasR* (DH164) expressing P*opdH*-GFP-*lacZ* promoter fusion at the *att*::Tn7 site | This study |
| PA14 Δ*lasR*Δ*tctE* P*opdH*-GFP-*lacZ* | | PA14 Δ*lasR*Δ*tctE* (DH) expressing P*opdH*-GFP-*lacZ* promoter fusion at the *att*::Tn7 site | This study |
| PA14 Δ*lasR*Δ*tctD* P*opdH*-GFP-*lacZ* | | PA14 Δ*lasR*Δ*tctD* (DH) expressing P*opdH*-GFP-*lacZ* promoter fusion at the *att*::Tn7 site | This study |
| PA14 Δ*lasR*Δ*tctED* P*opdH*-GFP-*lacZ* | | PA14 Δ*lasR*Δ*tctED* (DH) expressing P*opdH*-GFP-*lacZ* promoter fusion at the *att*::Tn7 site | This study |
| PA14 att::P*tac*-mKate | | PA14 WT with two tandem copies of mKate2 under a synthetic *tac* promoter integrated at the Tn7 *att* site | This study |

| | | | |
|---|---|---|---|
| PA14 Δ*lasR*Δ*rhlR* P*rhlI-lacZ* | DH3309 | PA14 Δ*lasR*Δ*rhlR* (DH2944) expressing P*rhlI-lacZ* promoter fusion at the *att*::Tn7 site | (7) |
| PA14 Δ*lasR*Δ*tctED* P*rhlI-lacZ* | | PA14 Δ*lasR*Δ*tctED* (DH) expressing P*rhlI-lacZ* promoter fusion at the *att*::Tn7 site | This study |
| PA14 Δ*lasR*Δ*opdH* P*rhlI-lacZ* | | PA14 Δ*lasR*Δ*opdH* (DH) expressing P*rhlI-lacZ* promoter fusion at the *att*::Tn7 site | This study |
| PA14 Δ*lasR*Δ*opdH* | | PA14 Δ*lasR (*DH164) with in-frame deletion of *opdH* | This study |
| PA14 Δ*lasR*Δ*opdH* + *opdH* | | PA14 Δ*lasR*Δ*opdH (*DH) with complementation of *opdH* at the native locus | This study |
| PA14 Δ*lasR*Δ*tctABC* | | PA14 Δ*lasR (*DH164) with in-frame deletion of *tctABC* | This study |
| PA14 WT *att::lacZ* | DH22 | PA14 WT with constitutive expression of *lacZ* | Roberto Kolter (8, 9) |
| PA14 Δ*lasR*Δ*betAB* | | In-frame deletion of *lasR* in Δ*betAB* (DH) | This study |
| PA14 Δ*lasR*Δ*betAB*Δ*tctED* | | In-frame deletion of *tctED* in Δ*lasR*Δ*betAB* (DH) | This study |
| PA14 Δ*tctD* | | PA14 WT (DH122) with in-frame deletion of *tct*D | This study |
| PA14 Δ*tctED* | | PA14 WT (DH122) with in-frame deletion of *tctE*D | This study |
| PA14 Δ*opdH* | | PA14 WT (DH122) with in-frame deletion of *opdH* | This study |
| PA14 Δ*tctABC* | | PA14 WT (DH122) with in-frame deletion of *tctABC* | This study |
| *E. coli* | | | |

| | | | |
|---|---|---|---|
| S17 λpir | DH71 | Used as a conjugation partner for introducing pMQ30 and GH121-based plasmids. | |
| DH5a | DH51 | Used to store/replicate plasmids. | Invitrogen |
| Plasmids | | | |
| pMQ30 EV | DH962 | Allelic replacement vector for use in yeast cloning, $Gm^R$ | (10) |
| GH121 EV | DH2830 | For inserting sequences at the *att*::Tn7 site via allelic replacement; $Gm^R$ | (11) |
| GH121_P*rhlI-lacZ* | DH3314 | GFP-*lacZ* under control of the *rhlI* promoter, for integration at the *att*::Tn7 site; $Gm^R$ | (7) |
| GH121_P*opdH-lacZ* | | GFP-*lacZ* under control of the *opdH* promoter, for integration at the *att*::Tn7 site; $Gm^R$ | This study |
| p*lasR*_KO | DH133 | PA14 *lasR* in-frame deletion construct; $Gm^R$ | (2) |
| pMQ30_*crc*_KON | DH3511 | *crc* (PA14_70390) native locus complementation construct; $Gm^R$ | (4) |
| pEX18_*tctED*_KO | | PA14 *tctED* in-frame deletion construct; $Gm^R$ | (12) |
| pMQ30_*opdH*_KO | | *opdH* in-frame deletion construct; $Gm^R$ | This study |
| pMQ30_*opdH*_KON | | *opdH* complementation construct at native locus; $Gm^R$ | This study |
| pMQ30_*tctE*_KO | | *tctE* in-frame deletion construct; $Gm^R$ | This study |
| pMQ30_*tctD*_KO | | *tctD* in-frame deletion construct; $Gm^R$ | This study |
| pMQ30_*tctABC*_KO | | *tctABC* in-frame deletion construct; $Gm^R$ | This study |

1. Rahme LG, Stevens EJ, Wolfort SF, Shao J, Tompkins RG, Ausubel FM. 1995. Common virulence factors for bacterial pathogenicity in plants and animals. Science 268:1899-902.
2. Hogan DA, Vik A, Kolter R. 2004. A *Pseudomonas aeruginosa* quorum-sensing molecule influences *Candida albicans* morphology. Mol Microbiol 54:1212-23.
3. Smith EE, Buckley DG, Wu Z, Saenphimmachak C, Hoffman LR, D'Argenio DA, Miller SI, Ramsey BW, Speert DP, Moskowitz SM, Burns JL, Kaul R, Olson MV. 2006. Genetic adaptation by *Pseudomonas aeruginosa* to the airways of cystic fibrosis patients. Proc Natl Acad Sci U S A 103:8487-92.
4. Mould DL, Stevanovic M, Ashare A, Schultz D, Hogan DA. 2022. Metabolic basis for the evolution of a common pathogenic *Pseudomonas aeruginosa* variant. Elife 11.
5. Mould DL, Botelho NJ, Hogan DA. 2020. Intraspecies signaling between common variants of *Pseudomonas aeruginosa* increases production of quorum-sensing-controlled virulence factors. mBio 11.
6. Dietrich LE, Price-Whelan A, Petersen A, Whiteley M, Newman DK. 2006. The phenazine pyocyanin is a terminal signalling factor in the quorum sensing network of *Pseudomonas aeruginosa*. Mol Microbiol 61:1308-21.
7. Harty CE, Martins D, Doing G, Mould DL, Clay ME, Occhipinti P, Nguyen D, Hogan DA. 2019. Ethanol stimulates trehalose production through a SpoT-DksA-AlgU dependent pathway in *Pseudomonas aeruginosa*. Journal of Bacteriology doi:10.1128/jb.00794-18:JB.00794-18.
8. Wang Z, Xiong G, Lutz F. 1995. Site-specific integration of the phage phi CTX genome into the *Pseudomonas aeruginosa* chromosome: characterization of the functional integrase gene located close to and upstream of *attP*. Mol Gen Genet 246:72-9.
9. Choi KH, Schweizer HP. 2006. mini-Tn7 insertion in bacteria with single *att*Tn7 sites: example *Pseudomonas aeruginosa*. Nat Protoc 1:153-61.
10. Shanks RM, Caiazza NC, Hinsa SM, Toutain CM, O'Toole GA. 2006. *Saccharomyces cerevisiae*-based molecular tool kit for manipulation of genes from gram-negative bacteria. Appl Environ Microbiol 72:5027-36.
11. Heussler GE, Cady KC, Koeppen K, Bhuju S, Stanton BA, O’Toole GA. 2015. Clustered Regularly Interspaced Short Palindromic Repeat-Dependent, Biofilm-Specific Death of *Pseudomonas aeruginosa* Mediated by Increased Expression of Phage-Related Genes. mBio 6:e00129-15.
12. Zhang L, Fritsch M, Hammond L, Landreville R, Slatculescu C, Colavita A, Mah TF. 2013. Identification of genes involved in *Pseudomonas aeruginosa* biofilm-specific resistance to antibiotics. PLoS One 8:e61625.
